# Cortical-layer EEG-fMRI at 7T: experimental setup and analysis pipeline to elucidate generating mechanisms of alpha oscillations

**DOI:** 10.1101/2025.09.09.674189

**Authors:** Daniel C. Marsh, Rodika Sokoliuk, Kevin M. Aquino, Daisie O. Pakenham, Ross Wilson, Rosa Sanchez Panchuelo, Matthew J Brookes, Simon Hanslmayr, Stephen D. Mayhew, Susan T Francis, Karen J Mullinger

## Abstract

Alpha band (8-13Hz) electroencephalography (EEG) oscillations play a key role in cognition, but their generating mechanisms are still poorly understood. Most studies investigating laminar origins of alpha oscillations have been conducted on animals using invasive intracranial recordings. To relate these findings to human alpha generation, non-invasive techniques need to be developed. Layer functional Magnetic Resonance Imaging (fMRI) at ultra-high field (UHF, 7T) allows for the interrogation of brain responses across cortical depths and combined with simultaneous EEG, provides the opportunity to gain new insight into human alpha generation mechanisms.

This work establishes a framework to study the generating mechanisms of electrophysiological signals non-invasively in humans using simultaneous EEG layer-fMRI. Data were acquired on 10 participants during an eyes closed/eyes open paradigm. We showed that in 9/10 participants the quality of EEG and Blood Oxygenation Level Dependent (BOLD) fMRI data were sufficient to observe a significant negative correlation between EEG alpha power and the BOLD signal in visual cortex grey matter to the eyes open/eyes closed task.

“Deveining” was performed to overcome the increase in BOLD signal toward the pial surface due to draining veins, and the effects of each of the steps in the deveining analysis on the cortical depth profiles of the negative alpha-BOLD correlations studied. The largest effect was dependent on the exclusion of voxels in the tissue immediately surrounding veins. Following deveining, the cortical depth profiles showed the negative alpha-BOLD correlations were significantly weaker in the middle depths compared with deep and superficial depths. When a boxcar rather than EEG alpha power was used to model the task, this depth-dependence was not seen, suggesting this was specific to spontaneous alpha-power modulations.

In conclusion, we have established a method to non-invasively interrogate the origins of electrophysiological signals. Our alpha-BOLD depth profiles suggest the alpha signal to an eyes open-closed task is generated in superficial and deep layers suggesting top-down processes.

## 1. Introduction

Electroencephalography (EEG) provides rich information on brain activity characterised as neuronal oscillations across different frequency bands, with the most prominent oscillations from the human cortex being in the alpha (8–13 Hz) band. Early seminal findings in cognitive neuroscience showed that the power of alpha oscillations over the occipital cortex increased when people closed their eyes (Adrian & Matthews, 1934; Berger, 1929). Later studies revealed the involvement of alpha oscillations in cognitive processes like perception, attention and memory performance (Klimesch et al., 1994, 1999; Lenartowicz et al., 2019; Sokoliuk et al., 2019; Webster & Ro, 2020) whilst dysfunction in alpha modulation has been shown in pathologies such as ADHD and schizophrenia (Abrams & Taylor, 1979; Lenartowicz et al., 2018, 2019; Li et al., 2008). Although evidence suggests that alpha plays an important mechanistic role in human cognition, the origins and generating mechanisms of this brain rhythm remain unclear which impairs a full understanding of its influence.

To improve understanding of the generators of scalp EEG signals it is highly desirable to identify the layers of cortex from which they originate (Bollimunta et al., 2008, 2011; Buffalo et al., 2011; Maier et al., 2010). Neural signals measured in the middle cortical layer (Layer 4) have been proposed to relate to bottom-up processing mechanisms and the integral involvement of the thalamic projections in the generation of these signals (Felleman & Van Essen, 1991; Hughes et al., 2004; Lawrence et al., 2018; F. H. Lopes da Silva et al., 1980; Lopes Da Silva & Storm Van Leeuwen, 1977). Whilst those measured in superficial (Layers 1-3) and deep (Layers 5 and 6) layers are believed to relate to top-down processes and the involvement of higher cognitive areas of the cortex such as the dorsal attention network (DAN) (Buffalo et al., 2011; Hughes et al., 2004; F. Lopes da Silva, 1991; Scheeringa et al., 2016). However, the poor spatial resolution of scalp EEG (Burle et al., 2015) limits its ability to determine the cortical layer origin of alpha oscillations. Instead, invasive electrophysiological recordings in animals (e.g. (Maier et al., 2010; Saalmann et al., 2012)) have been used to determine information on a submillimetre scale. Therefore, the development of a non-invasive method to elucidate the origins of EEG signals in humans is highly desirable. Functional MRI (fMRI) provides sufficient spatial resolution to localise brain activity to cortical layers. In recent years, there has been a rapid growth in the number of human laminar fMRI studies (Norris & Polimeni, 2019) and availability of ultra-high field (UHF, 7T and above) MR scanners. UHF fMRI has facilitated the study of cortical layer responses at high spatial resolution due to the increased signal-to-noise ratio (SNR) and increased contrast-to-noise ratio of the Blood Oxygenation Level Dependent (BOLD) signal due to amplified susceptibility effects (van der Zwaag et al., 2009, 2016), making it the go-to field strength for layer fMRI studies (Huber, 2018).

At 1.5 and 3T, combined EEG-fMRI has provided deeper understanding of the neurophysiology of brain activity by analysing covariations in neural and BOLD signals (Deligianni et al., 2014; Mayhew & Bagshaw, 2017; Mullinger et al., 2017; Plis et al., 2018; Sabatinelli et al., 2006; Sadeh et al., 2010; Whittingstall et al., 2007) including the laminar fMRI correlates of electrophysiological signals (Scheeringa et al., 2016). EEG-fMRI at 7T has been shown to be possible (Abbasi et al., 2015; Brookes et al., 2008; Jorge et al., 2015; Meyer et al., 2020; Vasios et al., 2006) but challenging due primarily to the increase in EEG artefacts with MR field strength (Mullinger, et al., 2008a), but possesses unique advantages for understanding the origins of electrophysiological signals.This work aims to address the methodological challenges of combining EEG with 7T layer-fMRI to study the origin of alpha oscillations in response to an eyes-open/eyes-closed task known to induce large changes in posterior alpha power. Specific methods covered include how to: 1) perform the optimal practical set-up of EEG and MRI equipment at 7T using a 64-channel EEG system and two 16-channel high-density array MR receive coils; 2) collect combined EEG-fMRI data with sufficient high spatial resolution to study cortical layers by using a 3D gradient echo-echo planar imaging (GE-EPI) acquisition with sufficient Field of View (FoV) to image V1, V2 and V3 visual areas; 3) account for the resultant longer repetition time (TR) (∼4 s) of this image acquisition than the TR used for standard spatial resolution EEG-fMRI studies (∼2 s) (Andreou et al., 2017; Moeller et al., 2008)); 4) account for the draining vein effect which alters the specificity of the GE-BOLD signal in layer-specific fMRI (Koopmans et al., 2010; Polimeni et al., 2010) when investigating EEG-BOLD relationships. An analysis pipeline was developed to investigate alpha-BOLD correlations across cortical depths. This paper aims to establish a framework by which the cortical and sub-cortical circuit mechanisms of electrophysiological signals can be established non-invasively in humans. We focus here on the origins of alpha oscillations due to their robust modulation in healthy participants and important role in cognitive processes and clinical relevance.

## 2. Methods

### 2.1. 7T EEG-fMRI data acquisition

This study was conducted with approval of the local ethics committee and complied with the Code of Ethics of the World Medical Association (Declaration of Helsinki). Ten healthy, experienced fMRI participants (four female, mean age 28±5 yrs) took part in the study. Each participant gave written, informed consent before the start of the experiment. The core protocol was an eyes open/eyes closed paradigm during which simultaneous EEG-fMRI data was collected. In a separate scan session (without EEG), fMRI data were acquired to define visual functional areas V1, V2 and V3, and structural MRI to define the grey matter (GM) for layer analysis.

MRI data were acquired on a 7 T Phillips Achieva MR scanner (Phillips Medical Systems, Best, Netherlands) using a circular polarized volume transmit coil (Nova Medical, Wilmington, USA). All EEG data were recorded using an MR-compatible EEG cap (EasyCap, Herrsching, Germany) with 63 scalp electrodes following the extended international 10 – 20 system and an additional channel for recording the electrocardiograph (ECG). The reference electrode was positioned at FCz and the ground electrode at AFz. MRI-compatible BrainAmp MR-plus EEG amplifiers (Brain Products, Munich) and Brain Vision Recorder (Version 1.10) were used for data acquisition.

#### Eyes-open/eyes-closed paradigm

An eyes-open/eyes-closed block paradigm was chosen to modulate alpha power and BOLD signals. A brief (100 ms) vibrotactile stimulus was applied to the index finger using a piezoelectric stimulation device (Dancer Designs, UK) to cue participants to the periods in which to switch between eyes open and eyes closed. A central fixation cross was presented via a projector screen during the eyes open periods and a blank screen during eyes closed periods. The paradigm consisted of alternating 30 s periods of eyes-open and eyes-closed, repeated four times per fMRI run. Four or five runs were acquired per participant dependent upon participant comfort (5/5 participants had five/four runs, respectively).

#### Session 1: EEG-fMRI and associated structural MRI measures

MRI data were collected using two 16-channel high-density array surface receive coils (MR Coils, Netherlands) (see ***Fig. 1A***) positioned over the participant’s occipital cortex using the known relative position of the EEG electrodes to the individual anatomy of each participant. Coban Tape was used to hold the surface coils in position prior to the participant being placed in the scanner (see ***Fig. 1B***). ***Figure 1C*** shows the positioning of the participant’s head, surface receive coils and EEG cap within the volume transmit coil. The use of surface receive coils rather than the standard 32-channel Nova head receive coil provided more space around the participant’s head, making EEG setup easier as well as providing higher image signal-to-noise ratio from the visual cortex and the capability to perform greater parallel imaging acceleration to speed up the image acquisition.

**Figure 1:**
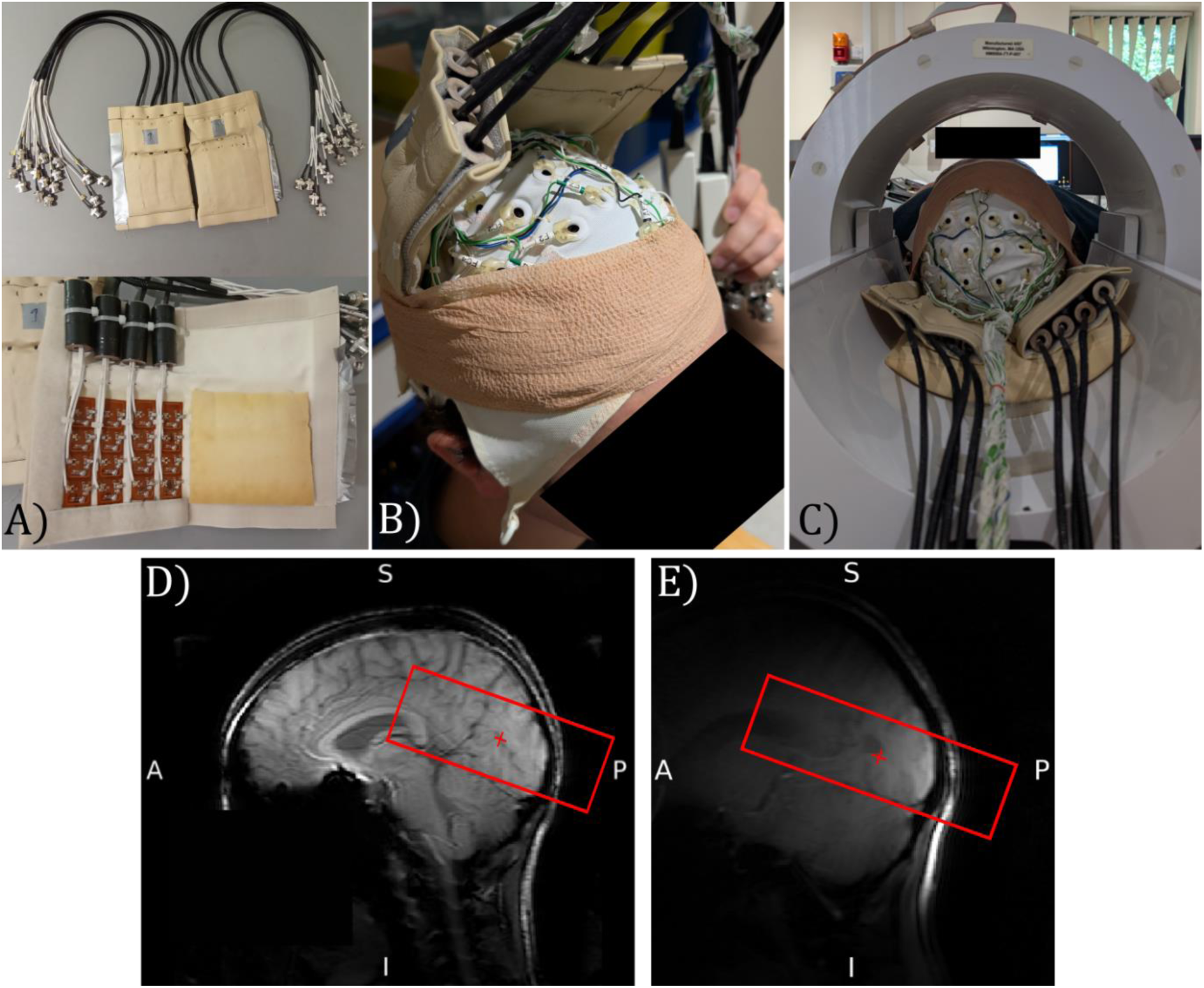
EEG-fMRI data acquisition. a) Two surface receive coils with 16 receive elements shown b) 64-channel EEG cap and position of the two surface receive coils on a participant held in place using Coban Tape prior to scanning, c) Position of the participant within the volume MR transmit coil with the EEG cap and surface receive coils in place, d) Survey image collected using the circular polarised MR transmit coil for both transmit and receive, and e) Survey image collected using the surface receive coils to localise sensitivity to brain regions. Red box shows the coverage of the imaging volume for the 3D GE-EPI functional data. Photographs depict an author with their consent for inclusion.

To ensure accurate positioning of the surface receive coils, a 3D survey scan was initially collected using the circular polarised transmit coil for both transmit and receive to provide full head coverage (***Fig. 1D***). A survey scan was then collected using the surface receive coils to localise the brain region under the coils (***Fig. 1E***). If required, the head was then moved relative to the surface receive coils to ensure optimal sensitivity in V1, V2 and V3 visual areas.

##### fMRI data

To achieve a 0.8 mm isotropic resolution, a 3D GE-EPI T2*-weighted sequence comprising 44 axial slices centred along the calcarine sulcus was collected with image-based (IB) shimming (TE/TR = 32/76 ms, FA = 28°, TRvolume = 3.8 s, Bandwidth (BW) in Measurement (M) = 740.4 Hz and BW in phase encoding (P) = 16.5 Hz, acquired matrix 88×168 (M x P in anterior-posterior (AP) and right-left (RL) respectively) (reconstructed matrix 256×256), SENSE 3.45×1.7 (P × S direction, the high density surface receive coils allowing a high SENSE factor to be preferentially used in the RL phase encoding direction). Each fMRI run comprised 68 volumes, resulting in a scan duration of ∼4 minutes. Cardiac and respiratory traces were recorded throughout each run using the scanner’s vector cardiogram (VCG) and respiratory belt.

##### EEG data

EEG data were acquired at a sampling rate of 5 kHz using hardware filtering of 0.016 – 250 Hz with a roll-off of 30 dB/octave at high frequency. Electrode impedances were kept below 20 kΩ. EEG artefacts from the MRI environment were minimised by isolating the EEG amplifiers from the scanner bed and reducing the MR room noise by switching off the cold head pumps during fMRI acquisition (Mullinger, et al., 2008a). Optimal correction of the gradient artefact was achieved by modifying the standard vendor 3D GE-EPI sequence to allow the TR to be set to 76ms to 6 decimal places ensuring it was equal to a multiple of the EEG sampling period, and by synchronising the EEG and MRI scanner clocks (Mullinger, et al., 2008b).

##### Field-map and Anatomical data

Prior to the fMRI acquisition, a B0 field-map (TR = 20 ms, TE/ΔTE = 5.92/1 ms, FA = 25°, SENSE 2, 4 mm isotropic, 64×64 matrix, 40 slices) was acquired with the same central position and angulation and image-based (IB) shim volume as the 3D-EPI fMRI data acquisition to allow this to be used for field-map based distortion correction of the 3D-EPI fMRI datasets. Following fMRI acquisition, magnitude and phase images of a phase-sensitive inversion recovery (PSIR) sequence (Mougin et al., 2016) were collected with the same central positioning and angulation as the 3D GE-EPI fMRI using the surface coils (0.7 mm isotropic, 252×250 matrix (M x P), 175 slices, TI1/TI2 = 780/2280 ms, SENSE: 2.2 x 2 (P x S direction)).

After completion of the EEG-fMRI scan session, the locations of the EEG electrodes on the scalp surface, and the shape of the participant’s head were digitally recorded using a Polhemus isotrack 3D system (Polhemus, Vermont, USA).

#### Session 2: Retinotopic Mapping and whole head Anatomical data

On a separate day, fMRI data (with no EEG) were collected using a 32-channel whole head receive coil (Nova-Medical) during retinotopic mapping (Wandell et al., 2007). Our standard 2D GE-EPI fMRI data acquisition protocol for retinotopic mapping with greater spatial coverage comprising 32 coronal oblique slices covering the entire visual stream (V1 to intraparietal sulcus (IPS)) and shorter TR (TE = 25 ms, 85° FA, TR = 2 s, 1.5 mm isotropic resolution, 124×121 matrix, SENSE 2.5 in foot-head P direction) with IB shimming was acquired. Eccentricity and polar angle maps were acquired using standard retinotopic mapping procedures comprising an expanding annulus and rotating wedges (Gardner et al., 2008), with 120 volumes collected per run. 4 runs were acquired in total, 2 for eccentricity (expanding and contracting) and 2 for polar angle (clockwise and anti-clockwise). A whole head T1-weighted anatomical PSIR (0.7 mm isotropic) was acquired with matched image acquisition parameters to those used in Session 1.

### 2.2. Analysis of EEG-fMRI data

#### EEG data

*Preprocessing*: EEG data were gradient artefact (GA) and pulse artefact (PA) corrected using Brain Vision Analyzer2 with the 3D-EPI volume acquisition markers and R-peaks from the VCG trace used to form sliding window templates (51 repetitions for GA template and 21 repetitions for PA template). Data were then filtered at 0.1 – 40 Hz, down-sampled to 500 Hz and exported to FieldTrip (Oostenveld et al., 2011) where all further analysis was performed. Data were epoched into segments, corresponding to time windows of TR/4, and visually inspected to identify noisy data, primarily due to movement, which were marked as noisy segments but remained in the data at this stage. On the continuous data, noisy channels were removed and Independent Component Analysis (ICA) performed on the remaining data, to remove components corresponding to eye blinks or horizontal eye movements (Joyce et al., 2003). The remaining ICs were back projected to channel space and data were re-referenced to an average of all the non-noisy channels. These continuous data were then filtered into the alpha (8 – 13 Hz) frequency band.

*Source-level analysis*: This involved three steps:

i. *<u>Construction of individual head models</u>*: Individual, four-layer boundary element (BEM) head models—comprising scalp, skull, cerebrospinal fluid (CSF), and brain—were constructed from each individual’s T1-weighted anatomical images using the ‘dipoli’ method implemented in the FieldTrip toolbox (Oostenveld et al., 2011). Using the fiducial points and head shape, individual electrode positions were aligned to the individual’s scalp surface derived from the T1-weighted scan.
ii. *<u>Identifying peak source location</u>:* Noisy time periods in the continuous EEG data (identified in the preprocessing, see above) were removed for this analysis stage, and the total duration of data for eyes-open and eyes closed conditions matched. All data were demeaned by subtracting the average over the whole time period for each electrode separately. Source localisation was then performed using a Linear Constraint Minimum Variance (LCMV) beamformer, with regularisation set to 5% of the maximum in the covariance matrix. Pseudo-T-statistic (Ŧ-stat) maps showing significant task-related changes in alpha-power between eyes-open and eyes-closed periods were calculated over the whole head (active window (eyes-open): 0.5-29.5s, passive window (eyes-closed): 30.5-59.5s). The peak location of the maximum change in alpha power between eyes-open and eyes-closed in the occipital cortex was identified from each participant’s Ŧ-stat map and chosen as the site of the Virtual Electrode (VE) - this ensured the maximum signal-to-noise ratio of the alpha-timecourse as the beamformer acts as a spatial filter (Brookes et al., 2008).
iii. *<u>Extraction of VE-timecourse</u>*: A VE-timecourse was extracted from the continuous pre-processed EEG data (see above) for each run (using a covariance matrix generated from all clean EEG data), and time locked to the start of the fMRI data acquisition. Noisy segments identified during pre-preprocessing were replaced by either the average over adjacent data segments (within a window of 4 segments if at least 2 individual segments within the window were artefact-free), else by the average over the whole eyes-open or eyes-closed condition for the run. The final alpha activity timecourse was demeaned and a Hilbert transform was applied to produce a continuous alpha power timecourse. To inspect final EEG data quality, a Fast Fourier Transform (FFT) of the VE timecourse (sampled at 500 Hz) for the 30s eyes-open and 30s eyes-closed periods, concatenated over blocks within a run, was taken for each run and averaged over runs. To assess whether there was an overall mean alpha power difference between the eyes-open and eyes-closed conditions, the average alpha power was calculated from the VE timecourses during each period and averaged across all runs for each participant. A paired Student’s t-test was performed to assess for a significant difference in alpha power between eyes-open and eyes-closed.

#### 3D-EPI fMRI data

##### Preprocessing

fMRI data were corrected for respiratory and cardiac physiological noise using retrospective image correction (RETROICOR) (H. Glover et al., 2000), and geometric spatial distortions using B0 field-map correction (FUGUE, FSL (https://fsl.fmrib.ox.ac.uk)) to provide good alignment of the fMRI with PSIR anatomical data. Motion correction of each fMRI run was performed using linear registration with MCFLIRT (Motion Correction using FMRIB’s Linear Image Registration Tool, FSL) and between runs using FLIRT (FMRIB’s Linear Image Registration Tool, FSL), with the transformation matrices calculated for each. These matrices were concatenated (convert_xfm, FSL) in a single transformation and applied to the fMRI data using 3^rd^ order spline interpolation to minimise spatial blurring (Polimeni et al., 2018). The result was that each fMRI volume was aligned to the central volume of the entire session.

##### EEG-fMRI General Linear Model (GLM)

Three GLMs were formed to allow us to optimise the analysis pipeline and subsequently establish if differences existed between the laminar origins of BOLD signal changes driven by the (i) mean effect of the task and (ii) volume-to-volume alpha power variability.

For GLM1 “Alpha”: the EEG alpha power VE timecourse was convolved with the FSL double gamma haemodynamic response function (HRF) and down-sampled to the sampling rate of the 3D-EPI fMRI data (1/ TRvolume =1/3.8 s =0.26 Hz) (resultant regressors are shown in ***Supplementary Information, Figure S1***). For GLM2 “Boxcar”: a boxcar task regressor with amplitude of “1” denoting eyes closed and “0” denoting eyes open periods was input into the GLM, instead of the alpha regressor used in GLM 1. For GLM3 “Alpha Orthogonalised”: the alpha power regressor was orthogonalized with the boxcar regressor, to remove the mean task effect from the alpha power variance. Each of these regressors were entered into first-level GLMs along with 6 motion parameters as regressors of no interest (FSL, FEAT v6.00).

Since alpha oscillations are known to correlate negatively with the BOLD response (alpha power increasing on eyes-closed) (Moosmann et al., 2003), negative beta weights corresponding to negative GLM z-statistics were taken forward for further analyses.

For each participant and GLM, a z-statistic map of the BOLD-alpha power correlation was calculated across all runs using a second-level fixed-effects analysis (threshold of z < −2.3, cluster correction (p < 0.05)(Nichols & Hayasaka, 2003)). Voxels with a large negative z-statistic (z < −2.3) indicate high alpha-modulation to the task. z-statistic maps were then up-sampled to 0.175 mm isotropic resolution using nearest neighbour resampling (FSL) to ensure voxels within each GM cortical depth for layer analysis (see ***Section 2.5***).

##### Anatomical data

For the PSIR images collected in Sessions 1 and 2, magnitude and phase were combined (Mougin et al., 2016) to form field-bias corrected partial and whole head PSIR images, respectively.

### 2.3. Defining visual boundaries from retinotopic mapping

Retinotopic mapping 2D-EPI fMRI data acquired in Session 2 provided functional V1, V2, and V3 boundaries to define regions of interest (ROIs) for EEG-fMRI analysis. Each of the four retinotopic runs were motion corrected and co-registered to the second run (MCFLIRT & FLIRT, FSL). The 0.7 mm isotropic whole head PSIR image was down-sampled to 1 mm^3^ and input into FreeSurfer (https://surfer.nmr.mgh.harvard.edu) (Dale et al., 1999; van der Kouwe et al., 2002) to compute GM, white matter (WM) and CSF surfaces. The 1 mm^3^ PSIR and functional retinotopic data were then aligned in mrTools (using mrAlign https://gru.stanford.edu/doku.php/mrtools/overview) (Gardner et al., 2018) before performing travelling wave analysis (Engel, 2012) to produce retinotopic phase maps on the cortical surface. These phase maps were used to manually define V1, V2 and V3 boundaries and functional ROIs, which were compared against the Benson Atlas ROIs (Benson & Winawer, 2018). For Participant 6, the Benson Atlas ROIs alone were used as the retinotopic phase maps were of insufficient quality to define visual boundaries.

To align the V1, V2 and V3 ROIs with the 3D EPI fMRI data, the Session 2 whole head 1 mm^3^ PSIR image was registered (6 degrees of freedom, FSL, FLIRT) to the Session 1 partial head PSIR image. The resultant transformation matrix was used to move V1, V2, and V3 ROIs into native 3D EPI fMRI space. ROIs were threshold to counteract smoothing due to the registration. Manual correction through visual inspection in FSLeyes (Smith et al., 2004) was applied to delete any overlap between the V1, V2, V3 ROIs resulting from the registration process. These final V1, V2 and V3 ROIs were then up-sampled to 0.175 mm for use in layer analysis.

### 2.4. Anatomical processing to generate a grey matter mask

Standard brain extraction tools (e.g. FSL’s Brain Extraction Tool (BET), and reconall in Freesurfer) are optimised for to segment whole brain images collected with a volume coil, so an optimal method was developed for our partial-head surface coil data.

First, the PSIR TI2 = 2280 ms magnitude image (which has the most GM/CSF contrast of two inversion times) was skull stripped using BET2 (FSL) (Smith et al., 2004). This image then underwent two erosion iterations (removing one voxel from the outer brain surface per iteration). This resulted in the best automated estimate of the brain which was then manually corrected to give an accurate definition of the individual’s brain. This corrected image was used to generate a binarised brain mask which was applied to skull strip the partial-head field-bias corrected PSIR image (which had the best GM/WM contrast for further analysis). This PSIR image was then segmented (FSL, FMRIB’s Automated Segmentation Tool (FAST) (Smith et al., 2004)), to obtain a GM binary mask. The skull stripped partial-head PSIR and GM binary mask were then matched to the field of view and geometry of the up-sampled (0.175 mm isotropic) 3D GE-EPI fMRI data. The GM mask was then manually corrected for any imperfections in the FAST segmentation of the GM ribbon such as merged gyri and veins within the GM.

### 2.5. Defining cortical depths and columns

The corrected GM mask image was labelled with values of CSF equal to 1, WM equal to 2, and GM equal to 3. This labelled GM mask was input to the LayNii software to define six cortical depths using the equivolume method (LayNii v2.0.0 (Huber et al., 2020)). Cortical columns were generated using the midpoint of the modelled depths (***Fig. S2***). The effect of the number of columns modelled using LN2_COLUMNS (LayNii) was explored for: i) 4000 columns, diameter ∼2 mm, and ii) 20000 columns, diameter ∼0.8 mm and described in ***Supplementary Information Section S2.1*** and ***Figure S3***.

Those columns to be included in the layer analysis were determined as illustrated in ***Figure 2***. A column was included if a voxel with significant negative activation (z < −2.3), reflecting alpha power-BOLD correlation, was contained at any depth of the whole column; a column was excluded if there was no significant negative voxel at any depth. This column inclusion was chosen because all the voxels across cortical depths are influenced by the draining vein effect (Turner, 2002; De Martino et al., 2013; Polimeni et al., 2010). This method is distinct from previous studies that include only voxels with z < −2.3 (Aitken et al., 2020; Haarsma et al., 2022).

**Figure 2:**
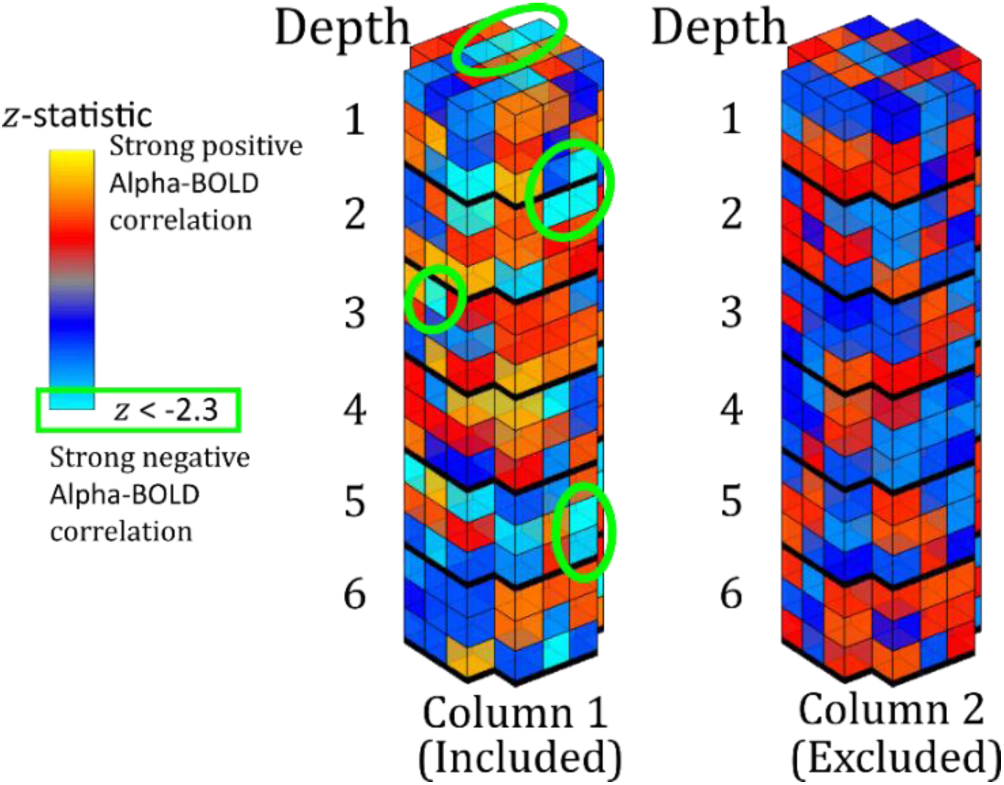
Schematic showing column inclusion. Column 1 contains multiple voxels with a significant negative alpha-BOLD correlation (z < −2.3 shown in light blue) and so the whole column is included. Column 2 has no significant negative alpha-BOLD correlation and is excluded.

For the included columns, those voxels to retain were evaluated based on β-weights. A high negative β-weight corresponds to voxels with a strong negative alpha-BOLD correlation; very low magnitude β-weights reflect voxels that do not respond to the task and which will propagate noise to other depths through deveining, see ***Supplementary Information Section S3***. This evaluation, led to the choice of retaining β-weights above a threshold of 5 % (of the absolute β-weights across all included columns) for subsequent analyses as this removes voxels dominated by noise whilst retaining those containing signal of interest.

### 2.6. Deveining optimisation

This process was all carried out only using the GLM1: Alpha only EEG regressor. The β-weights in each column were corrected for the draining vein effect using spatial deconvolution deveining (Havlicek & Uludağ, 2020; Heinzle et al., 2016; Markuerkiaga et al., 2021; Marquardt et al., 2018, 2020) which assumes that each layer contains a mixture of neural signals from that specific layer along with signals from all preceding deeper layers (Havlicek & Uludağ, 2020; Heinzle et al., 2016; Markuerkiaga et al., 2016). This spatial signal subtraction involves correcting the current layer in the column with a weighted sum across the preceding layers. An estimate of the voxel-wise venous cerebral blood volume (CBVv) required for spatial deconvolution deveining was estimated by calculating the voxel-wise amplitude of low frequency fluctuations (ALFF) (Guidi et al., 2020) from the mean variance of the BOLD signal during all eyes-closed periods for each participant. The following parameters within the spatial deconvolution deveining model were assessed for their impact on cortical depth profile shape.

#### 2.6.1. Effect of the scaling parameter λ

When implementing spatial deconvolution deveining (LayNii v2.0.0) a scaling parameter, λ, is used together with the ALFF to estimate the cerebral blood flow (CBF) (Huber et al., 2020) between tissue layers (Markuerkiaga et al., 2016). We assessed the effect of the physiologically range of λ of 0.2, 0.25, and 0.3 (Markuerkiaga et al., 2016) on deveining.

#### 2.6.2. Effect of ‘Global Mean’ versus ‘Column Profile Mean’ for cortical depth profile estimation

Cortical depth profiles were calculated in each visual region (V1, V2 and V3) using the ‘Global Mean’ and ‘Column Profile Mean’ method. The ‘Global Mean’ method calculated the mean β-weight for each depth across all columns. The ‘Column Profile Mean’ calculated an individual cortical depth profile for each column and then averaged these column profiles - this is likely more physiologically accurate but also more susceptible to noise, as a single depth within a column containing only a few active voxels will contribute equally to another column containing all possible active voxels (∼ 350 voxels) within that depth. The effect of column diameter on the ‘Global Mean’ method is explored in the ***Supplementary Information S2.1***.

#### 2.6.3. Effect of the proximity to veins

Draining veins impact layer-dependent GE-BOLD both within the veins as well as surrounding voxels (Fracasso et al., 2021). Thus, despite the exclusion of veins by applying a stringent GM mask, cortical depth profiles are likely to be affected by proximity to draining veins. After calculating the ‘uncorrected’ and ‘deveined’ cortical depth profile for each column in V1, the gradient of the profiles was calculated by fitting to a 1^st^ order polynomial (MATLAB). The relationship between this gradient and the column proximity to veins was assessed by creating a vein mask in V1 from the mean 3D GE-EPI fMRI image. First, the mean 3D GE-EPI image was spatially smoothed using a Gaussian 2 mm full-width-half-maximum kernel, the smoothed image was then subtracted from the original unsmoothed mean 3D GE-EPI image, and an upper threshold manually applied to create a binary ‘vein mask’ (***Fig. S5***). The vein mask was restricted to V1 using the V1 functional ROI. A ‘surrounding vein mask’ was then generated, by dilating the ‘vein mask’ four-fold and subtracting the original undilated vein mask producing a ∼0.8 mm width ring.

For each participant, the change in the gradient of β-weights due to deveining was computed in each of the remaining columns. To determine whether a spatial dependence of the gradient change existed, the columns were classified into those with the lower 25%, middle 50% and upper 25% of gradient profile changes. The percentage overlap of these three classifications with the ‘surrounding vein mask’ was calculated. A repeated measures ANOVA between each classification was used to determine if there was a significant difference between spatial patterns of the three classifications.

### 2.7. Cortical depth profiles calculation

The ‘uncorrected’ and the ‘deveined’ cortical depth profile in V1, V2 and V3 ROIs for those voxels with a negative β-weight (representing the areas of negative alpha-BOLD correlation) was calculated for each participant using only the columns containing significant (z<-2.3) voxels, as described in Section 2.5. This was computed for both the ‘uncorrected’ and ‘deveined’ data, since the deveining process can alter the sign of the β-weight resulting in different voxels being used in the profile estimation.

Firstly, for each participant in each of the visual ROIs (V1, V2 and V3), the cortical depth profile before and after deveining were normalised to the range of β-weights in their profile. A weighted average of the normalised cortical depth profiles across ROIs was then computed based on the number of columns present in each ROI for each participant and the group mean and standard error in cortical depth profile was computed. This was only carried out for the layer profiles from the Alpha EEG regressor GLM (see ***Section 2.2***)

Once columns with the largest gradient changes had been removed (see ***Section 2.6.3***), cortical depth profiles for ‘uncorrected’ and ‘deveined’ data were assessed for all three GLMs ([i] Alpha EEG regressor, [ii] Boxcar regressor, [iii] Alpha orthogonalized with boxcar regressor), using the same methods described above. This allowed us to test if the optimal pipeline could differentiate between the laminar origins of BOLD signal changes driven by the (i) mean effect of the task and (ii) alpha power variability.

### 2.8. Statistical analysis

Layer β-weight profiles after deveining were interrogated for statistically significant (p<0.05) differences across cortical depths for the four sets of cortical depth profiles outlined in ***Section 2.7*** using SPSS v29.0.1.0.

The cortical depth profiles with all columns included (i.e. not considering column proximity to veins) were evaluated using a one–way repeated measures ANOVA (rm-ANOVA) and if significant, post-hoc t-tests were performed to identify which depths were driving the overall effect and to localise peaks and troughs within the cortical depth profiles. Bonferroni correction was applied for multiple comparisons.

For the three sets of profiles where columns close to veins had been excluded, a two-way rm-ANOVA was applied across all depth profiles (factors: depth profile; GLM type [denoting the different regressors input into each GLM]). Significant differences in depth profile with GLM type were interrogated through the interaction of factors in the ANOVA (depth profile x GLM type). Then separate one–way repeated measures ANOVA (rm-ANOVA) were performed upon the cortical depth profiles from each GLM, to test for a significant effect of depth upon β-weight. If a significant effect was found, then post-hoc t-tests were performed to identify which depths were driving the overall effect and to localise peaks and troughs within the depth profiles. Bonferroni correction was applied to control for multiple comparisons.

## 3. Results

### 3.1. EEG and fMRI data quality

***Figure 3*** shows for the ten participants the EEG alpha power time course, normalised to the maximum value, for the eyes-open/eyes-closed paradigm recorded during fMRI. For most participants and runs, a periodic pattern of alpha power increasing during eyes-closed is seen. For Participant 7, run 1 was excluded from further analyses due to the signal in the final ∼80 s being replaced with the mean alpha value. When considering the average alpha power, there was the expected significant (p<0.05, students paired t-test) reduction during eyes open compared to eyes closed across the group, (see Table S1 for mean EEG alpha power values for each participant). ***Figure 4*** shows the average power spectrum of the VE time course during eyes-closed and eyes-open periods across all runs, normalised to the maximum alpha (8 - 13Hz) band power during the eyes-closed period. This highlights the variation in alpha power amplitude between participants, most participants show a visible difference in alpha power during eyes-closed compared to eyes-open.

**Figure 3:**
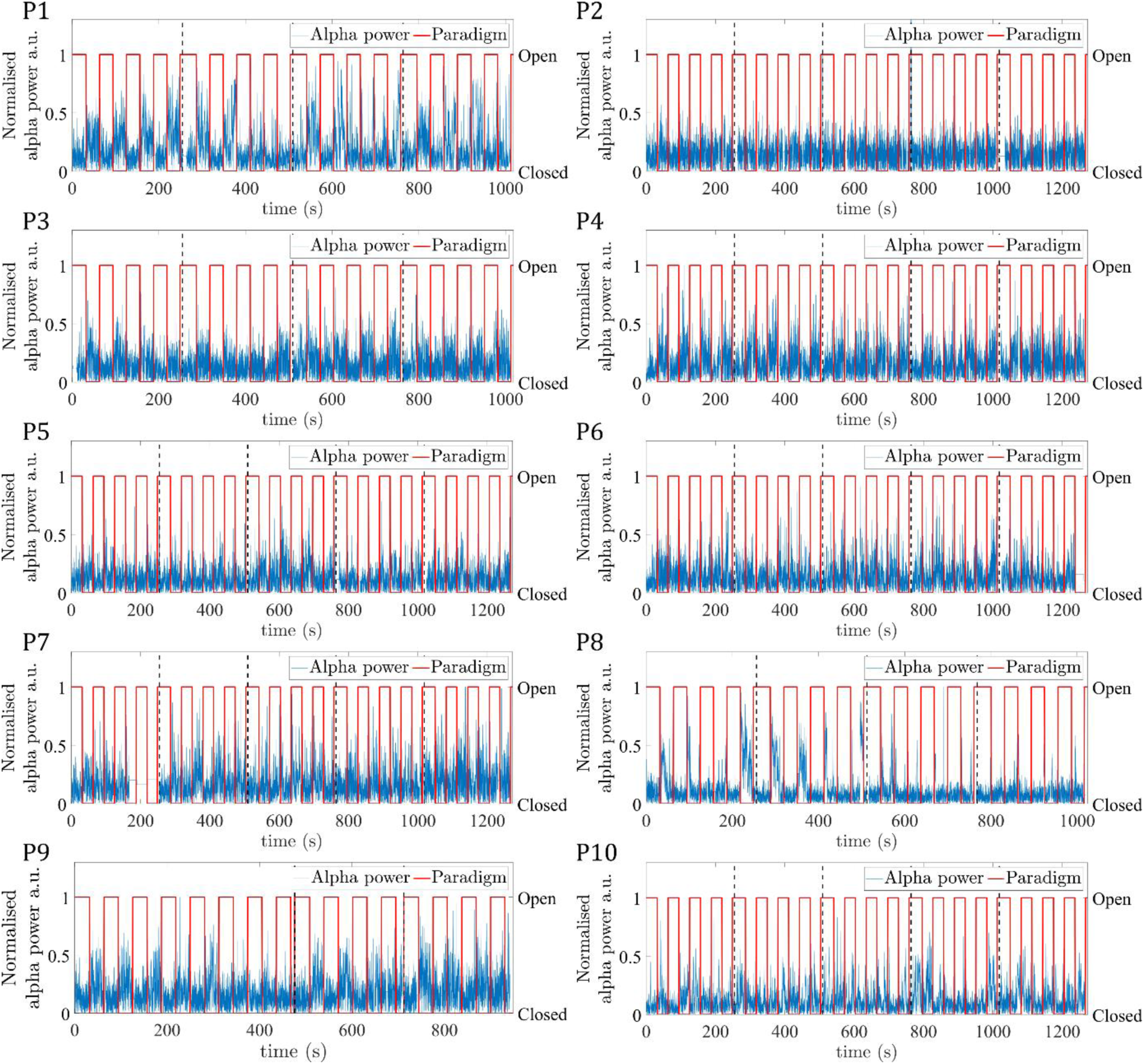
EEG alpha power VE time course (blue) at the peak alpha power location concatenated across all runs (indicated by vertical black dashed lines), for the 10 participants (P1-P10). Paradigm timing is overlaid (red), with periods of eyes-open/eyes-closed denoted by a value of 1/0.

**Figure 4:**
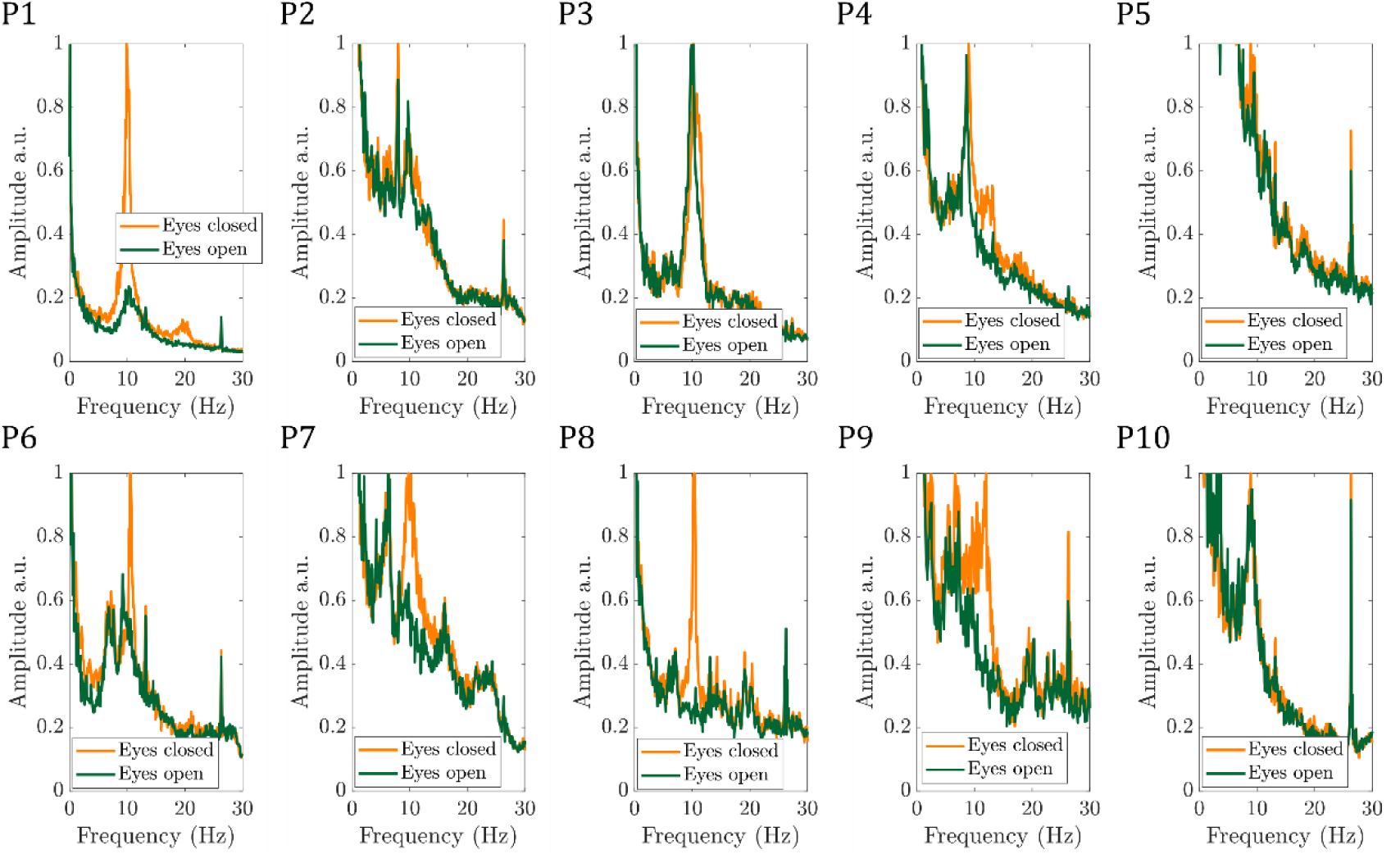
Average power spectrum of the VE timecourse during the eyes-open (green) and eyes-closed (yellow) for the 10 participants (P1-P10), normalised to the maximum spectrum power in the alpha (8 - 13Hz) band during eyes-closed.

***Figure 5*** shows the mean 3D-EPI image for each participant after pre-processing and the corresponding temporal signal-to-noise (tSNR) map, no signal loss or distortions due to the presence of the EEG electrodes are seen. The mean tSNR within the V1-3 ROIs showed good consistency across participants at 10.9±0.8 (mean±SEM), with the surface receive coils resulting in higher tSNR nearer the cortical surface than white matter.

**Figure 5:**
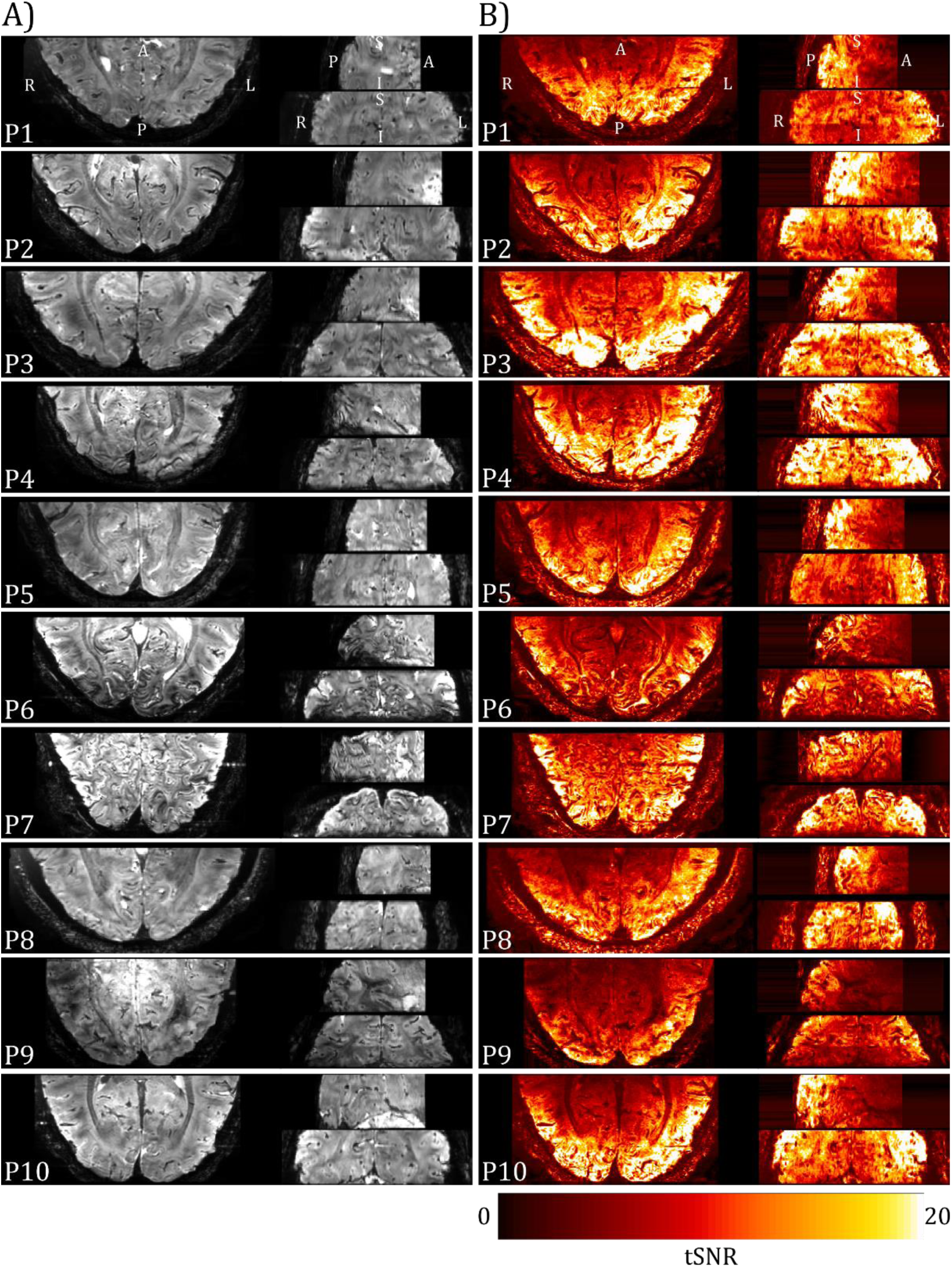
(A) Mean 3D-EPI images for each participant collected using the two 16-channel high-density array surface coils with the EEG cap in place (axial left, sagittal upper right, coronal lower right) following pre-processing (motion and distortion correction). (B) Corresponding temporal SNR (tSNR) images.

### 3.2. fMRI response to alpha power modulations and depth definition

All results, unless stated otherwise, are derived from the alpha GLM not orthogonalized with the task boxcar. Individual participant z-statistic maps of the alpha-BOLD correlation during the eyes-open/closed task are shown in ***Figure 6***, associated EEG alpha power time course regressors are shown in ***Fig. S1***. All participants showed significant negative correlation in the primary visual cortex that was spatially specific to GM. Close correspondence of the activation maps with the PSIR GM mask for each participant is shown, with a mean overlap of 53±8% for all participants (see ***Table S2***), highlighting good registration between the datasets.

**Figure 6:**
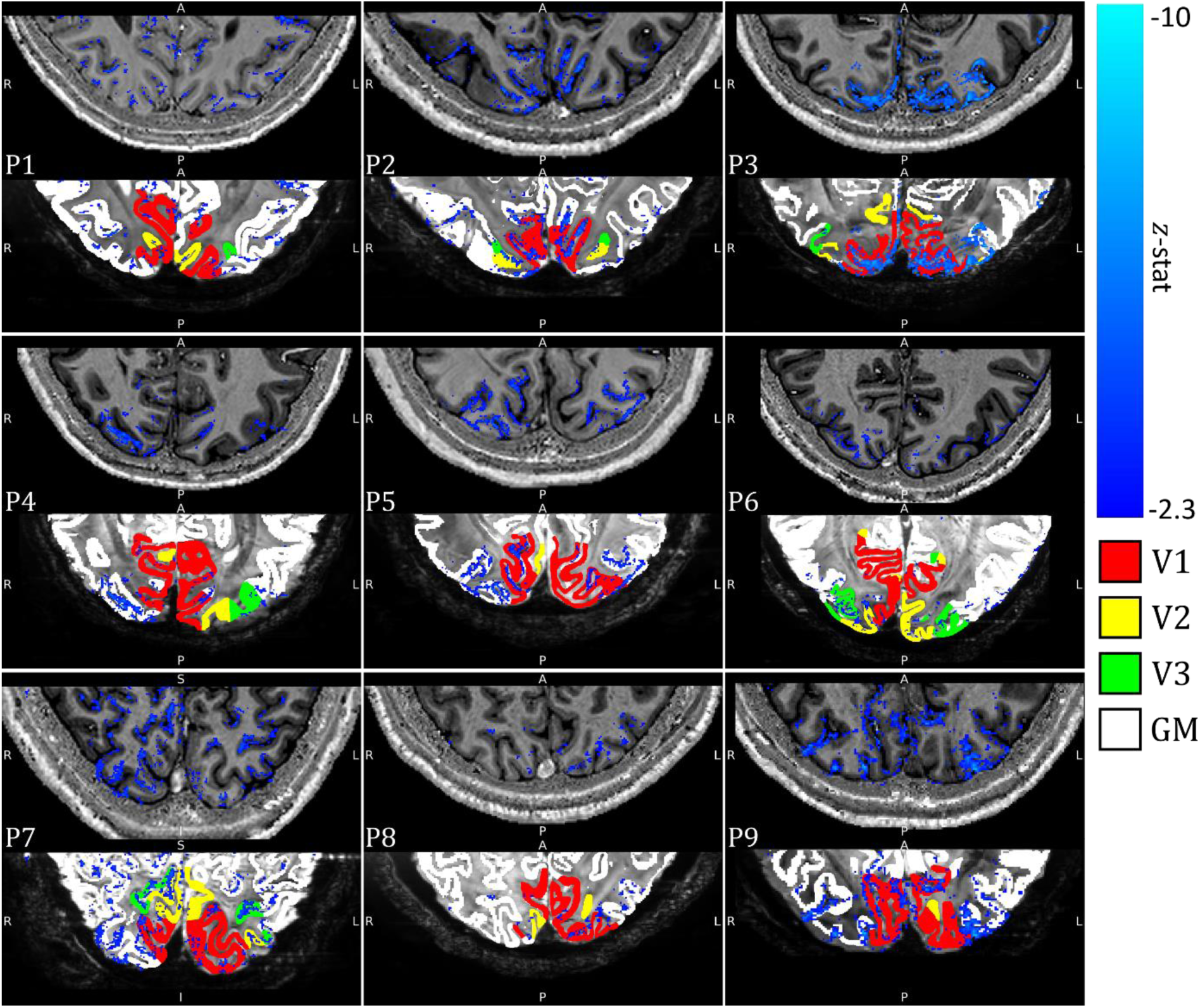
Negative contrast z-statistic map (threshold at z < −2.3 and cluster corrected p < 0.05) showing regions of significant alpha-BOLD correlation overlaid on each participant’s PSIR (Top panel). Threshold negative contrast z-statistic map superimposed on the mean 3D-EPI, final GM mask used for cortical depth modelling (white) with visual areas V1 (red), V2 (yellow), and V3 (green) shown (Bottom panel). Results shown for a single slice for each participant. See Figure S7 for a comparison of z-statistic maps for each of the GLMs used to model the BOLD response.

Participants 2, 3, 4, 5, 6, 7 and 9 showed a greater number of activated voxels than 1 and 8 (see ***Table S2***). Participant 10 showed a lack of alpha power modulation between eyes-open and eyes-closed (***Fig. 3&4***) and no significant negative alpha-BOLD correlation within the primary visual cortex (see ***Fig. S6***) so was excluded from further analyses.

### 3.3. Impact of analysis steps on cortical depth profile

The tissue segmentation and upsampling to yield six equivolume modelled depths and the cortical columns is shown in ***Figure S2*** for each participant. The effect of deveining parameters and constraints on the visual cortex cortical depth profile using 20000 columns is outlined in this section. Results are predominately shown for Participant 1, but all participants are shown for steps where effects were variable across participants.

#### 3.3.1. Effect of β-weight thresholding

***Figure 7*** shows the cortical depth profile after deveining of the normalised β-weight with ‘no threshold’ and a ‘5% threshold’ (***Supplementary Information Section S3***). The cortical depth profiles are very similar, a finding seen for the majority of participants (data not shown). A ‘10% threshold’ was deemed too stringent (see ***Fig. S4***), and so a 5% threshold was used for all further analyses.

**Figure 7:**
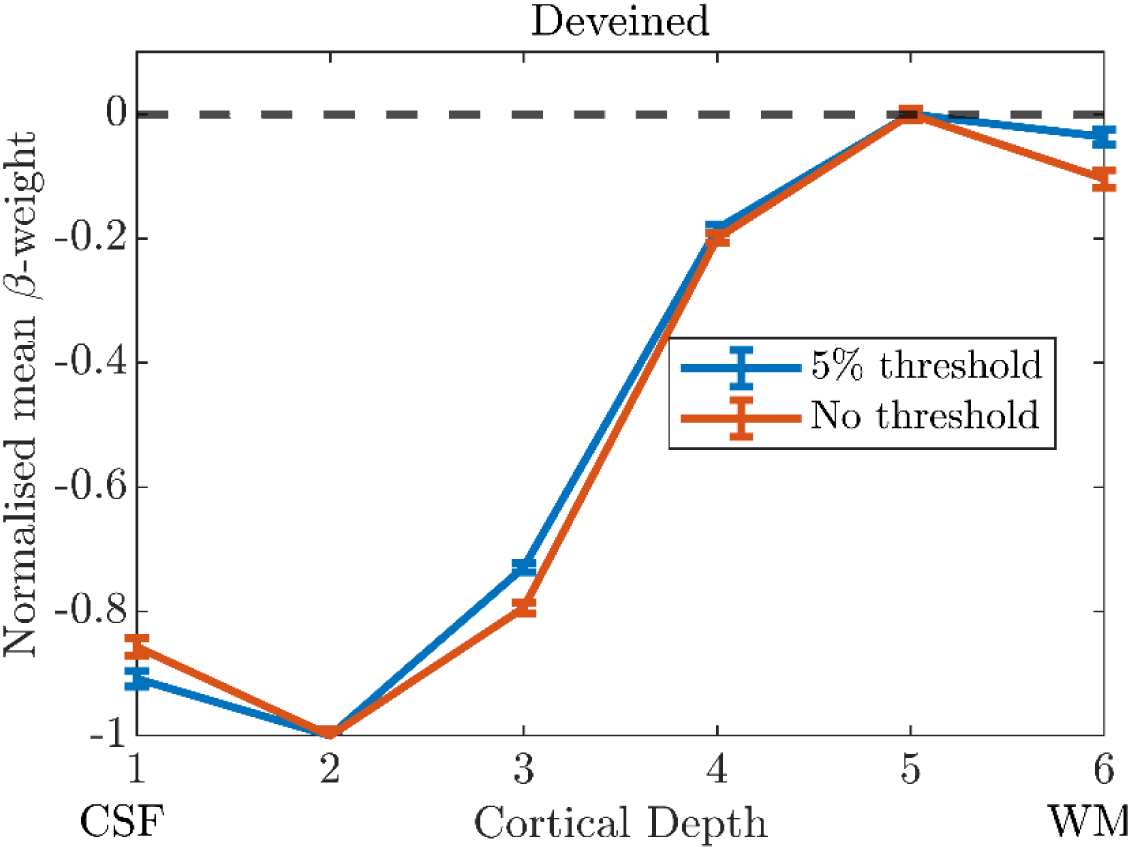
The ‘deveined’ cortical depth profiles of normalised β-weights from V1 using no threshold (all β-weights, orange) and a threshold of > 5% of maximum β-weight (blue). Data shown for 20,000 columns and the ‘Global Mean’ method. Error bars show standard error on the mean. Results shown for Participant 1.

#### 3.3.2. Choice of the scaling parameter, λ, in spatial deconvolution deveining

***Figure 8*** shows cortical depth profiles for λ of 0.2, 0.25, and 0.3. Before normalisation (***Fig. 8A***), for all participants the same mean β-weight is observed at depth 6 and diverges towards superficial depths, since λ scales the level of contribution from deeper layers subtracted at each depth. After range normalisation (***Fig. 8B***), each λ value resulted in a very similar cortical depth profile shape, and so the default value of λ of 0.25 was used for all further analyses.

**Figure 8:**
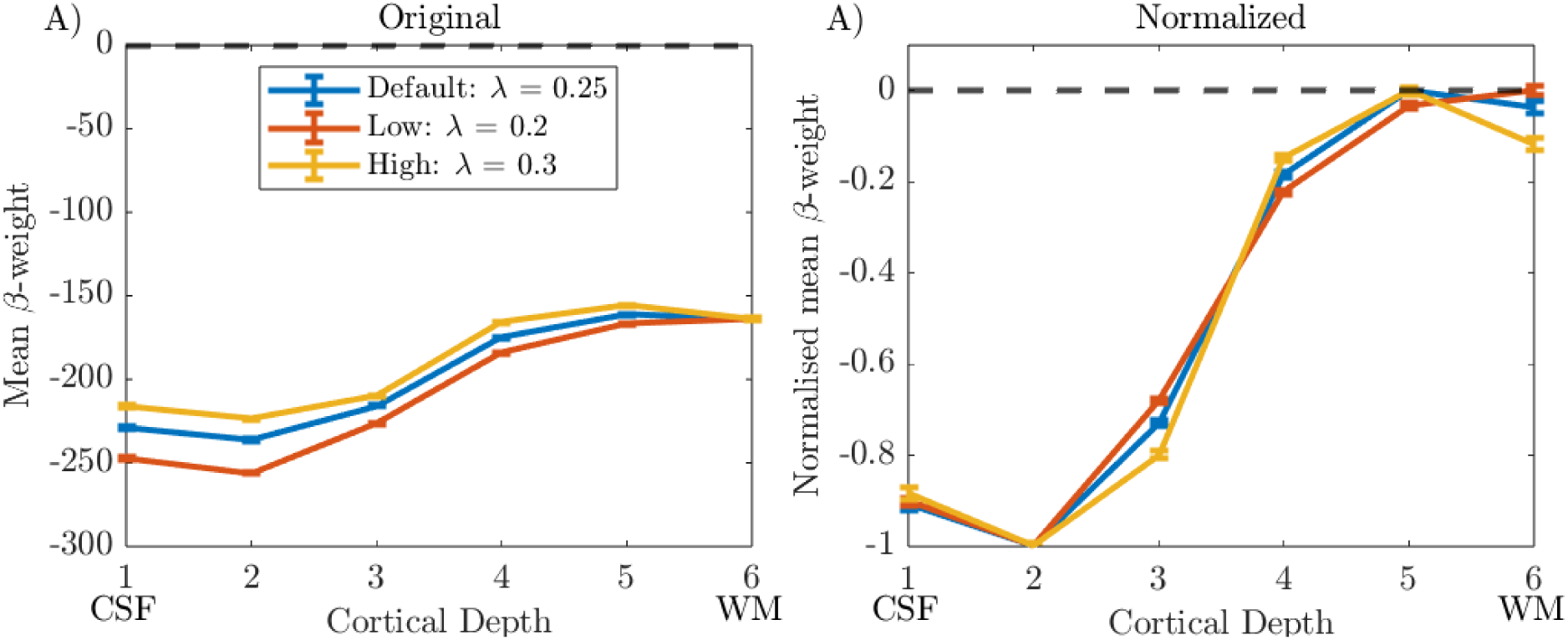
‘Deveined’ cortical depth profiles from V1 using 20,000 columns and the ‘Global Mean’ method shown for data deconvolution with default (λ = 0.25), low (λ = 0.2) and high (λ = 0.3) λ values in LayNii. Error bars show standard error on the mean. Data shown for Participant 1.

#### 3.3.3. ‘Global Mean’ versus ‘Column Profile Mean’ cortical depth profiles

For all participants, very similar cortical depth profiles were observed for the ‘Global Mean’ and ‘Column Profile Mean’ approaches to averaging β-weights for both the ‘Uncorrected’ (***Fig. 9A.i***) and ‘Deveined’ (***Fig. 9A.ii***) data. Therefore, the ‘Global Mean’ method was used for the group analysis as it is less prone to noise.

**Figure 9:**
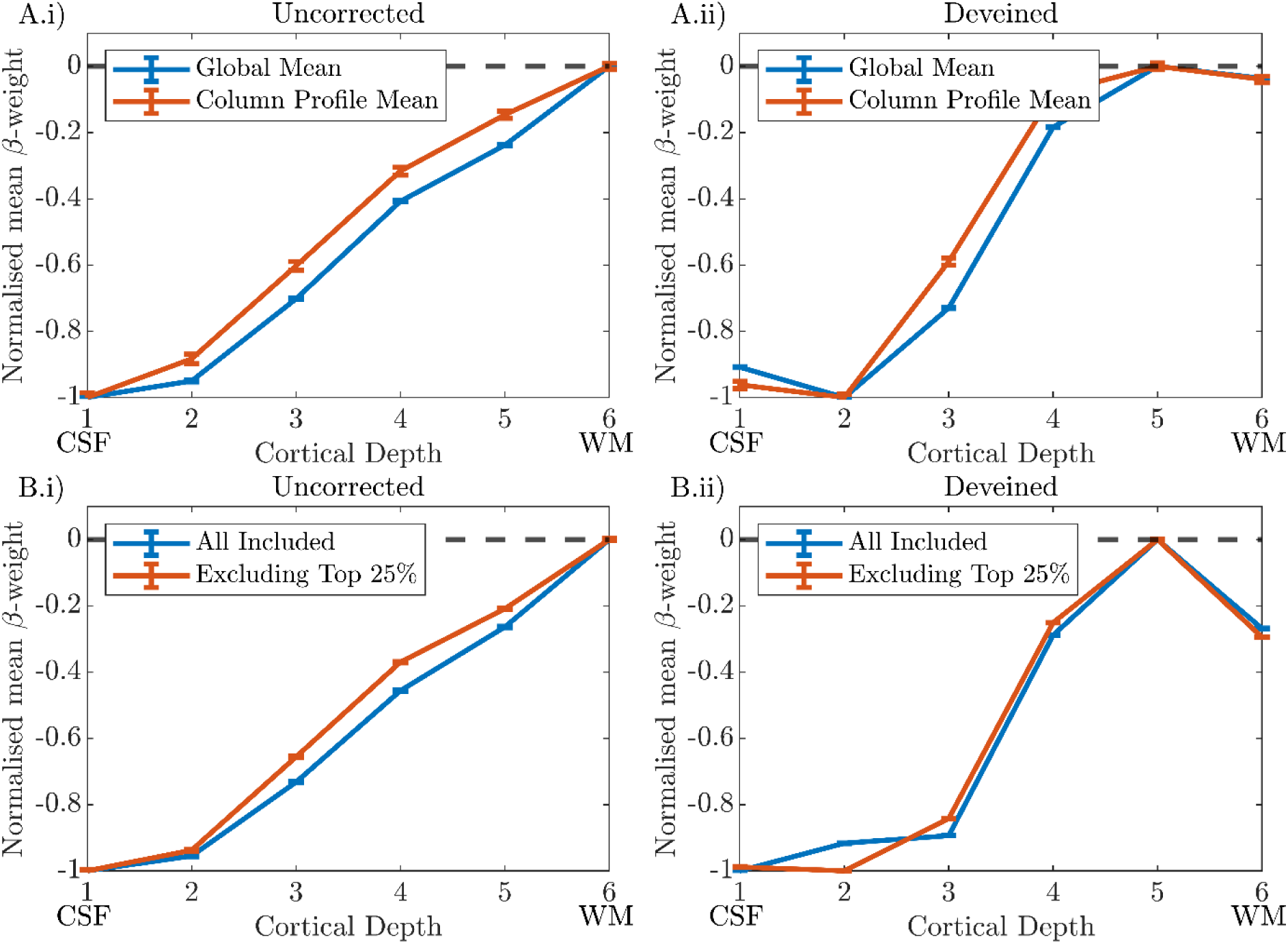
Cortical depth profiles from V1 for a single participant (P1) shown for ‘Uncorrected’ (i) and ‘Deveined’ (ii) GE-BOLD data. **Row A)** Blue and Orange lines indicate the ‘Global Mean’ and ‘Column Profile Mean’ methods respectively. **Row B)** Blue indicates the profile including all available columns, Orange shows the profile resulting from excluding the columns with the highest change in profile gradient (top 25%) due to deveining. All plots: error bars show the standard error on the mean (SEM) for the mean β-weight at each cortical depth.

#### 3.3.4. Effect of proximity to veins on cortical depth profile

In some columns, the sign of the gradient of the cortical depth profile switched after deveining, suggesting over correction across depths (see ***Fig. S8***). We hypothesised this was due to the effect of draining veins on tissue (Fracasso et al., 2021) surrounding the vein mask.

To test this hypothesis, all active voxels were ranked by the magnitude of their gradient change (of the column in which they sat) after deveining and overlap with the ‘vein mask’ and ‘surrounding vein mask’ was assessed. We observed no significant (p>0.05, rm-ANOVA) spatial relationship between the gradient change voxels and the ‘vein mask’. Across participants, 6.9 ± 1.0% of voxels classified in the upper 25% of cortical depth profile gradient changes overlapped with the ‘vein mask’, 4.0 ± 0.5% of voxels in the middle 50% gradient change voxels and 4.7 ± 0.9% of voxels in the lower 25% gradient change voxels (mean ± standard error over all participants. For the ‘surrounding vein mask’, 32.7 ± 3.0% (mean ± standard error over all participants) of voxels classified in the upper 25% of gradient changes overlapped, significantly greater (p<0.05, rm-ANOVA) than the 24.6 ± 2.8% overlap with the middle 50% gradient change voxels and 24.5 ± 3.1% overlap with the lower 25% gradient change voxels (p < 0.05, post-hoc t-tests).

The result of removing the columns in the top 25% of gradient changes and recalculating using the ‘Global Mean’ method is shown in ***Figure 9B*** for Participant 1, for whom this had a minimal effect. ***Figure 10*** shows this for all participants, showing that the deveined cortical depth profiles after range normalisation had considerable participant variability.

**Figure 10:**
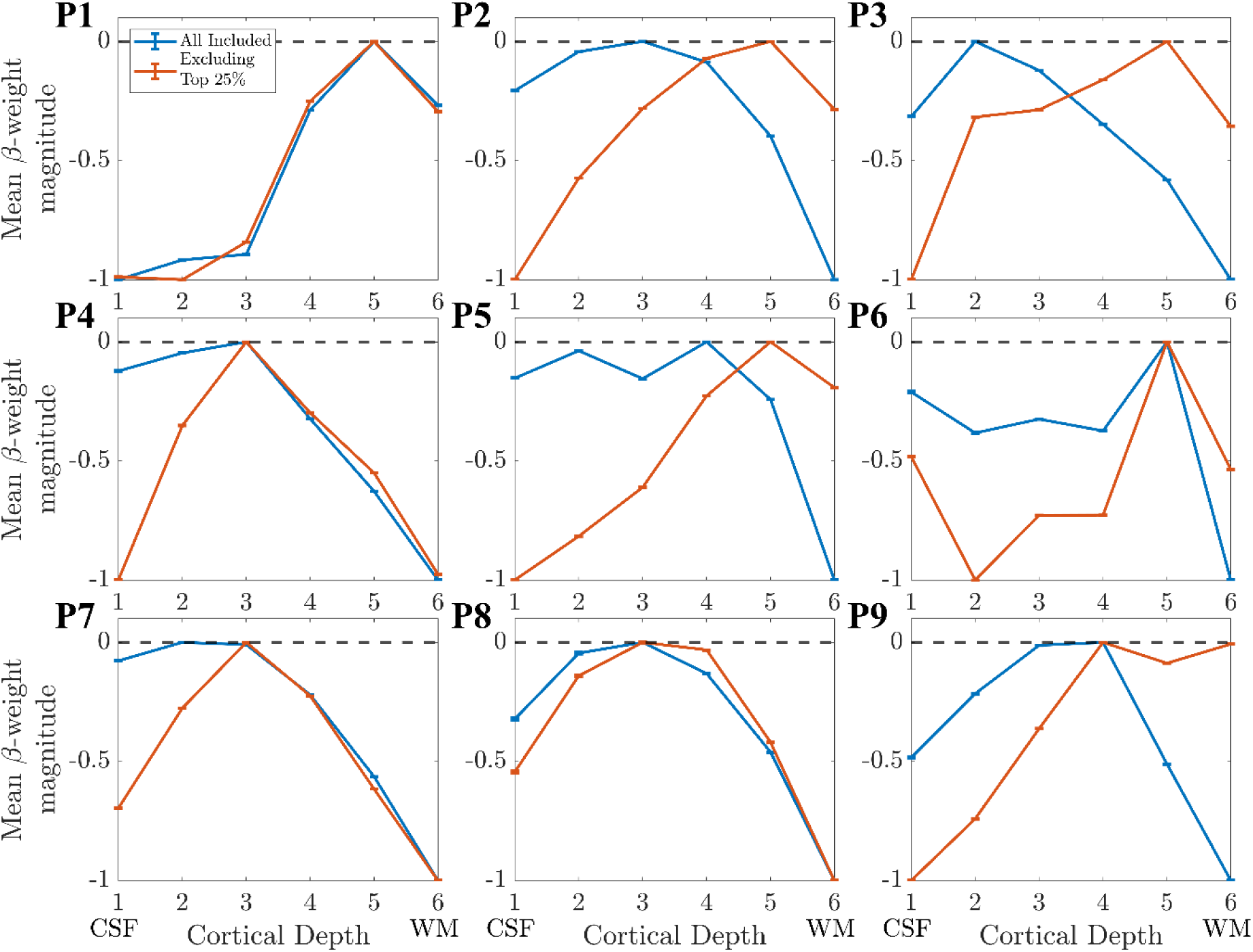
The layer profiles from V1 for participants 1-9 after ‘Deveining’ when 1) all available columns are used (orange) and 2) the columns with the highest change in profile gradient (top 25%) are excluded (blue).

Since it is shown in ***Supplementary Information S2.1*** that the profiles are not greatly affected by the size of the columns, a larger column size (smaller total number of columns) is preferable to maximise the presence of voxels in all layers which is more physiologically plausible. Therefore, remaining analysis was carried out using 4000 columns.

### 3.4. Assessing group cortical depth profiles

#### Alpha GLM profile

The group cortical depth profile for the alpha power GLM was first assessed using 4000 columns, a noise β threshold of 5% and λ = 0.25, and ‘global mean’ voxel averaging, and all columns containing voxels with <-2.3 included, see ***Figure 11***. The ‘uncorrected’ cortical depth profile showed across all participants a clear increase in β-weight magnitude towards the pial surface (***Fig. 11A***) driven primarily by the draining vein effect and therefore not of interest for understanding neuronal response origins. In comparison, the ‘deveined’ data showed an “n” shape profile across participants, with the weakest negative alpha-BOLD correlation in the middle cortical depths. A repeated measures ANOVA showed that the mean β-weight was significantly different (p = 0.005, rm_ANOVA) across the cortical depths with post-hoc t-tests revealing this was driven by: β-weights at cortical depth 1 less than depth 2 (p = 0.046), and β-weights at depth 6 less than depths 3, 4 and 5 (p = 0.031, p=0.002, p<0.001, respectively). There were no other significant differences between depths.

**Figure 11:**
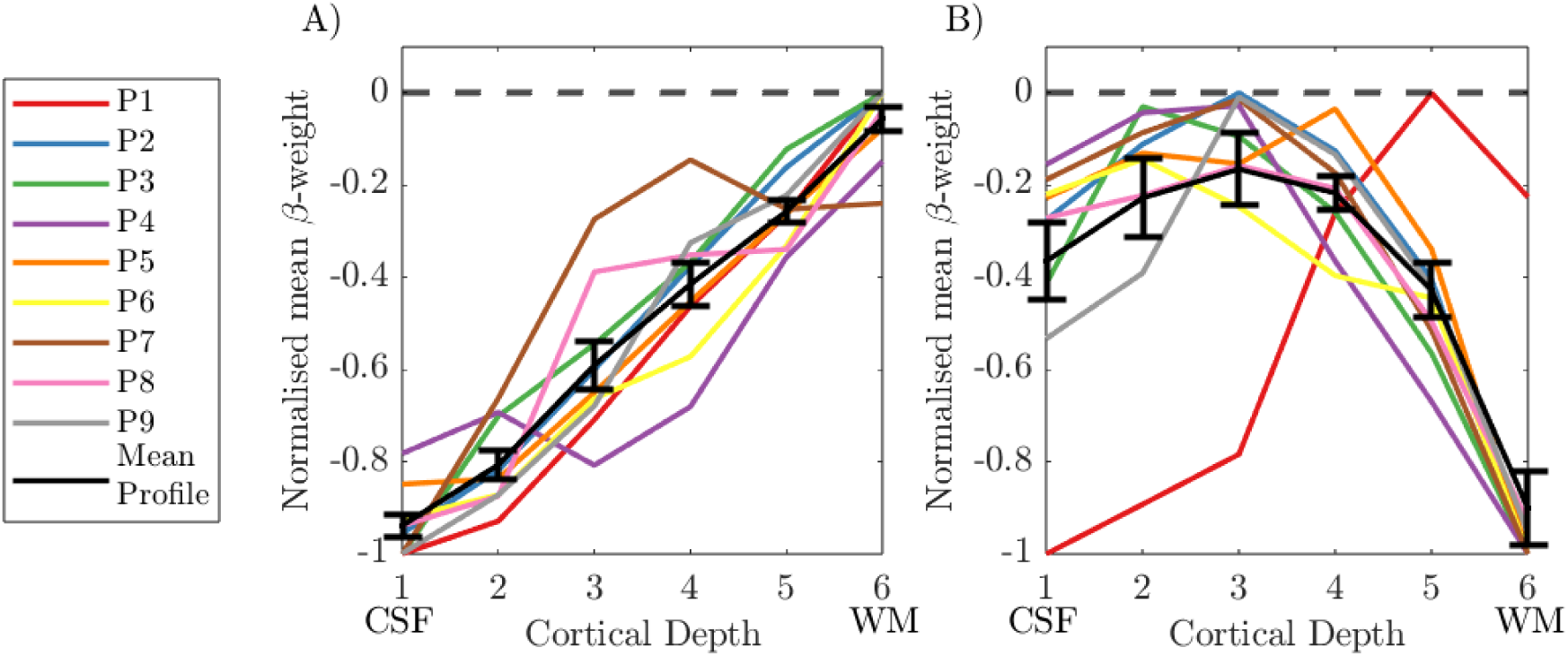
Cortical depth profiles showing the weighted average over V1–3 in normalised β-weights over cortical depths for (A) the uncorrected data and (B) deveined data. Individual participant profiles are shown with the group mean in black. Each participant’s response was normalised to the range of β-weights across depths for that participant. Error bars showing the standard error on the mean (SEM) over participants. CSF = cerebrospinal fluid; WM = white matter.

#### Comparison of depth profiles between GLM approaches

***Figure 12*** shows the ‘uncorrected’ and ‘deveined’ cortical depth profile for the GLMs: 1) Alpha power only, 2) Boxcar of task only and 3) Alpha power orthogonalized to boxcar; all calculated after excluding the columns where the gradient changed the most due to deveining (see ***Section 2.6.3***). In the ‘uncorrected data’ in all cases the typical bias to the cortical surface is seen (***Fig. 12* *column i***). After ‘deveining’ a greater variation in cortical depth profiles was seen across participants (***Fig. 12A.ii***) compared to no exclusion of columns (***Fig. 11B***). However, the exclusion of columns in close proximity to veins is likely to better represent the underlying neuronal activity, and so only this analysis is considered when comparing GLMs.

**Figure 12:**
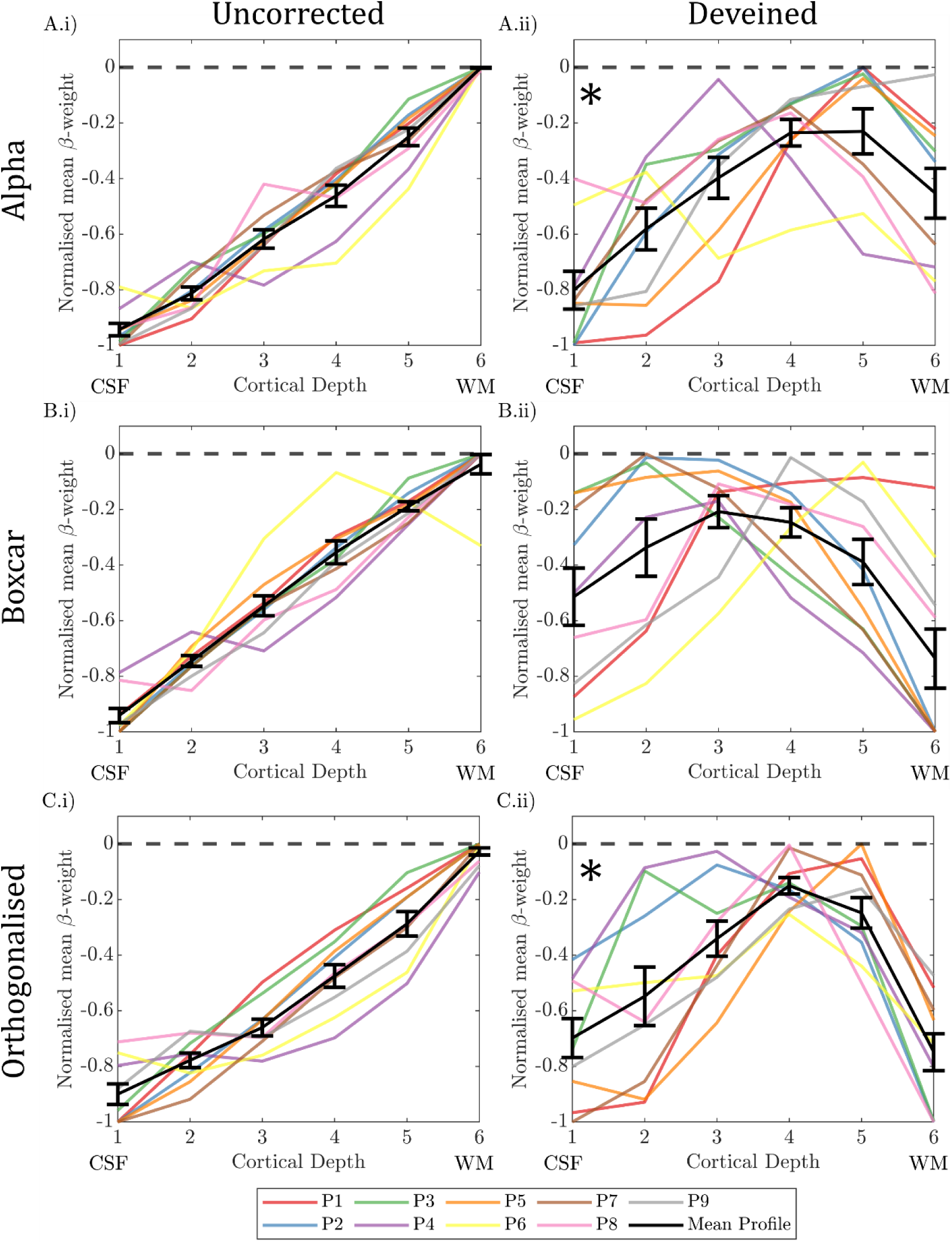
Cortical depth profiles showing the weighted average over V1–3 in normalised β-weights over cortical depth for (column i) uncorrected data and (column ii) the deveined data where columns likely to be close to veins were excluded (top 25% of gradient change columns [Section 2.6.3]). Normalised β-weights are calculated from three different GLMs featuring either A) alpha power regressor, B) boxcar regressor and C) alpha regressor orthogonalized with the boxcar regressor. Individual participant profiles are shown with the group mean in black. Each participant’s response was normalised to the range of β-weights across depths for that participant. Error bars showing the standard error on the mean (SEM) over participants. CSF = cerebrospinal fluid; WM = white matter. * denotes significant (p<0.017, Bonferroni corrected) effect of cortical depth on β-weights (one-way rm-ANOVA).

The deveined profiles across all GLMs revealed a significant effect of depth (p=0.004, two-way rm-ANOVA) and a trend for an interaction between depth and GLM type (p=0.083, two-way rm-ANOVA). To understand what was driving this interaction, depth profiles for each GLM type were interrogated separately with a significance threshold level being p < 0.017 (accounting for Bonferroni correction). A significant “n” shaped cortical profile was shown for the alpha only regressor GLM (***Fig. 12Aii***, p = 0.007 rm-ANOVA), with post-hoc t-tests revealing this was driven by β-weights at cortical depth 1 less than depth 4 (p = 0.010), and β-weights at depth 6 less than depth 5 (p = 0.024), no other post-hoc pairwise comparisons showed significant differences. When the boxcar regressor GLM was interrogated, no significant change in normalised β-weights with cortical depth was found (***Fig 12Bii***, p = 0.069 rm-ANOVA). For alpha orthogonalized with the boxcar regressor, a significant “n” shaped cortical profile was observed (***Fig 12Cii***, p = 0.003 rm-ANOVA), with post-hoc t-tests revealing this was driven by β-weights at depth 1 less than depth 3 (p = 0.004) and 4 (p = 0.003), and β-weights at depth 6 being less than depth 4 (p = 0.001) and 5 (p < 0.001), no other post-hoc pairwise comparisons showed significant differences.

## 4. Discussion

This work presents the first 7T EEG-fMRI layer study to investigate the origins of electrophysiological brain activity on the laminar scale. We present an experimental set-up to perform simultaneous UHF EEG-layer-fMRI and an associated analysis pipeline for which we show the influence of parameter choice and ordering of analysis steps. Results show that alpha-BOLD correlations appear strongest in the deep and superficial layers of visual cortex, when alpha is modulated through an eyes open-closed paradigm.

### Experimental set-up for simultaneous EEG-fMRI at 7T

It is well recognised that there are challenges related to performing EEG-fMRI at 7T (Warbrick, 2022). These include the path of the EEG cabling (Meyer et al., 2020; Pan et al., 2023) and safety related to the magnitude of heating effects confounded due to increased specific absorption rate (SAR) at higher field (Mullinger, Debener, et al., 2008). Here, the use of two 16-channel surface receive coils inside the transmit coil provided sufficient space for the EEG cap and cabling without the need to adapt EEG hardware (Meyer et al., 2020) (***Fig. 1***). A 3D-EPI fMRI acquisition was chosen since it has higher image and temporal SNR than 2D-EPI at 0.8 mm isotropic resolution (Poser et al., 2010; Van Der Zwaag et al., 2011) and allows parallel acceleration in both phase-encoding dimensions making it more time efficient at sub-millimetre resolution to minimise the TRvolume. Importantly, the low Ernst flip angle of 3D-EPI minimised the SAR (0.5 W/kg) and thus safety concerns due to heating during the EEG-fMRI acquisition (Hawsawi et al., 2017; Mullinger, et al., 2008c). The 3D-EPI fMRI data timhad high tSNR with minimal B0 distortion or B1 dropout despite the closer proximity of the surface coils to the EEG electrodes (***Fig.5***). This experimental set-up combined with optimised post-processing methods to remove EEG artefacts (Allen et al., 1998, 2000; Brookes et al., 2008), ensured good quality EEG-fMRI data.

The EEG data quality in the alpha frequency band was good with all participants exhibiting the expected increase in average alpha power during eyes-closed compared to eyes-open (***Table S1***); as expected the degree of alpha power modulation varied between participants (***Figs 3* *and 4***) (Haegens et al., 2014; Nakanishi et al., 2020). High quality, source localised EEG alpha-power timecourses were extracted from occipital cortex and used as GLM regressors to model the BOLD fMRI data. A significant negative correlation between EEG alpha power and the BOLD signal was observed in 9/10 participants (***Figs 6* *and S6***) localised to the GM (***Table S2***). Together, our results showed that the data were of sufficient quality to investigate the layer dependence of EEG alpha signals, an important goal of EEG-fMRI at 7T (Meyer et al., 2020; Scheeringa & Fries, 2010).

### Localisation of BOLD responses and deveining

The overlap of voxels with significant EEG-alpha-BOLD correlation and the GM mask increased significantly when the GM mask was dilated by 2 voxels, with a greater increase when dilating the mask towards the CSF (***Table S2***). This demonstrates the expected blurring towards the pial surface due to draining veins.

A number of approaches have been proposed for correcting GE-BOLD data for the draining vein effect (Fracasso et al., 2018; Gau et al., 2020; Guidi et al., 2020; Havlicek & Uludağ, 2020; Heinzle et al., 2016; Kashyap et al., 2018; Markuerkiaga et al., 2016; Marquardt et al., 2018). Here we chose to use spatial deconvolution deveining (Havlicek & Uludağ, 2020; Markuerkiaga et al., 2021; Uludag & Havlicek, 2021), the most physiologically based approach, but also the newest. Thus the impact of deveining parameters on depth profiles was assessed.

We investigated the effect of voxels dominated by noise on deveining and showed that excluding voxels within columns whose β-weights were below 5% of the maximum β-weight provided a good compromise between removing those voxels in a column which were dominated by noise and not removing meaningful signal (***Fig 7*** and ***Section S3***). However, this exclusion did not greatly change the shape of the cortical depth profile. Similarly, changing λ, a scaling parameter for the contribution of the deep layers in the deveining, made little difference to the range-normalised profiles (***Fig. 8B***), in agreement with a previous study (Markuerkiaga et al., 2021). Performing averaging on a ‘Global Mean’ versus ‘Column Profile Mean’ to generate cortical depth profiles also had little effect on cortical depth profiles (***Fig. 9A***).

### Effect of column size on layer profile

We considered the effect of column size on depth profile (as previously explored by (Huber, 2020)) by varying the number of columns in the GM ROI. We found the cross-sectional area of the columns is important, reducing column area often resulted in the top and bottom depths of the column containing no voxels limiting the deveining process. In addition, columns must span the venous tangential branch length and be >0.6mm (Huber, 2020). However, if the column area is too large then the assumption that tissue within the column is affected by the same physiological and physical effects will not hold. We showed that the depth profile shape was not changed greatly by the number of columns (4000-20000) (***Fig. S3B***). Thus 4000 columns, corresponding to a column diameter of approximately 2 mm, was used to ensure voxels were present at all depths of all columns.

### Proximity of columns to veins

The largest effect on the depth profile of alpha-BOLD correlations was seen when accounting for the proximity of columns to veins (***Fig. 10***). Importantly, these were not columns within veins, as these had been initially excluded from the GM mask, but those surrounding the vein mask. Those columns for which the gradient of the depth profile changed sign after deveining were typically in the tissue surrounding large veins where the draining vein effect still dominated (Bause et al., 2020; Koopmans et al., 2010; Moerel et al., 2018; Polimeni et al., 2010), in contrast to capillaries and venules which the deveining algorithm is designed to correct. To our knowledge, the effect of proximity to veins for layer profiles has not been investigated previously. Our results suggest it is important to consider excluding a larger area of tissue around veins for layer profiles in future studies.

### Interpretation

The group cortical depth profile showed the expected draining vein effect, with the negative alpha-BOLD correlation showing strongest magnitude at the pial surface (***Figs 11A* *&* *12* *column i***). These profiles, however, are not meaningful in interpreting which neural assemblies are responding to drive the alpha signal due to the dominance of the draining vein effect (Koopmans et al., 2010; Polimeni et al., 2010). After ‘deveining’ the data, results showed a significant n-shaped profile of negative alpha BOLD correlation across cortical depths for all GLMs considered (***Fig. 12***).

Whilst it is impossible to infer alpha-generation from BOLD-alpha power layer correlations (Scheeringa et al., 2016), we showed that the n-shaped profile of the alpha-BOLD correlation across cortical depths appeared to be driven by the spontaneous alpha power modulations (***Fig. 12 Cii***) rather than task condition changes (eyes open/closed boxcar, ***Fig 12 Bii***). It is important to note that the maintenance of the n-shaped profile for the alpha orthogonalized with the boxcar regressor is despite a clear reduction in the number of voxels where a significant negative alpha-BOLD correlation was observed compared with the alpha only regressor (***Fig. S7 rows B&C***). Together our results suggest the cortical depth profiles relate specifically to alpha power. Our results show that the negative alpha-BOLD correlations are weakest in the middle cortical depths, suggesting layer 4 is not the generator of spontaneous alpha-power fluctuations during an eyes open/closed task. These findings juxtapose the well-established hypothesis that resting alpha is generated by bottom-up processing with thalamocortical feedforward alpha mostly terminating in layer 4 (Mumford, 1991; Thomson & Bannister, 2003). However, our result does align with the findings of van Kerkoerle *et al*. who showed that the alpha layer profile for a sustained visual stimulus was dominated by the deeper and superficial layers using invasive electrophysiology recordings in monkeys (Van Kerkoerle et al., 2014). As our paradigm comprised long 30 s periods of eyes-open or eyes-closed we could expect our alpha-BOLD layer profiles to exhibit a similar shape, driven by similar cortical processes, which perhaps differ from “resting state” alpha power fluctuations, when eyes are permanently closed. In addition, in several other invasive electrophysiology studies on monkeys, alpha sources have been reported in both superficial and deep layers, and in general have been suggested to be strongest in superficial cortical layers (Bollimunta et al., 2011; Buffalo et al., 2011; Haegens et al., 2015). However, these results come with the challenge of interpretating their dependence on the location of the reference electrode (Haegens et al., 2015). Taken together, the results we show in ***Figure 12*** appear most in line with findings of such invasive electrophysiology studies.

The only prior work using non-invasive simultaneous EEG-fMRI to study the estimated depth profile relationship between EEG activity and BOLD signal was at 3T study (Scheeringa et al., 2016). They presented attentional visual task. Responses to visual stimuli were strongest in the superficial layers, with significant alpha-BOLD correlations over all cortical depths, while attention modulation of alpha was only seen in superficial layers. However, Scheeringa *et al*. did not apply deveining and so results likely to be confounded by the draining vein effect as correlations were generally highest at the GM-CSF boundary, akin to ***Figures 11A* *&* *12* *column i***.

### Future work

Using the methodology developed here, it could be established if alpha generators vary between tasks (e.g. attention/working memory/passive visual tasks) to elucidate more about functional role of alpha power. In addition, an alternative approach for identifying the layer specific origins of electrophysiological signals is to combine EEG with a non-BOLD fMRI acquisition (unaffected by draining vein signals) such as Vascular Space Occupancy (VASO) (Huber et al., 2021). However, an EEG-VASO technique poses additional challenges from EEG-BOLD approach including safety considerations for EEG due to the higher SAR of VASO (Huber et al., 2021), inherently lower tSNR than BOLD and longer TR typically required. Another approach which is showing promise for layer profile responses is magnetoencephalography (MEG). However, currently this method has only differentiated electrophysiology signal between infragranular and superficial depths (Bonaiuto et al., 2018). OPM-MEG may allow profiles to be determined over more depths, getting closer to the signals from the underlying cytoarchitecture (Helbling, 2025). Overcoming technical challenges and pushing boundaries in both of these approaches are important future avenues to explore.

## Conclusion

This work provides experimental set-up and data analysis methods to study simultaneous EEG-fMRI at a laminar level. EEG data showed alpha modulations in response to the eyes-open/eyes-closed task and, when correlated with BOLD fMRI data, allowed layer specific alpha-BOLD associations to be studied. The resultant cortical depth profiles showed a significant significantly weaker magnitude of the negative alpha-BOLD correlation in middle layers of the cortex, implying that the alpha response to an eyes open/closed paradigm is generated through top-down neuronal mechanisms from the deeper and superficial layers.

## Acknowledgements

We thank Renzo Huber, Faruk Gulban for insightful conversations regarding deveining. This work was supported by a Leverhulme Trust Research Project [grant number RPG-2014-369]. Daniel C. Marsh, Susan T. Francis and Karen J. Mullinger are supported by National Institutes of Health Grant, USA [grant number R01MH128475]

## Ethics and Integrity

All data and code will be shared where possible, upon request.

Data collection was funded by the Leverhulme Trust (Research Project Grant to K.J.M. RPG-2014-369).

There are no conflicts of interest.

All data collection was approved by the University of Nottingham Medical School Research Ethics Committee. All volunteers gave written, informed consent.

## Notes

### Competing Interest Statement

The authors have declared no competing interest.

### Summary of Updates

Removal of duplicate final paragraph in section 1. Introduction

