## Supplementary Information for "Cortical-layer EEG-fMRI at 7T: experimental setup and analysis pipeline to elucidate generating mechanisms of alpha oscillations"

### S1. EEG-fMRI regressors

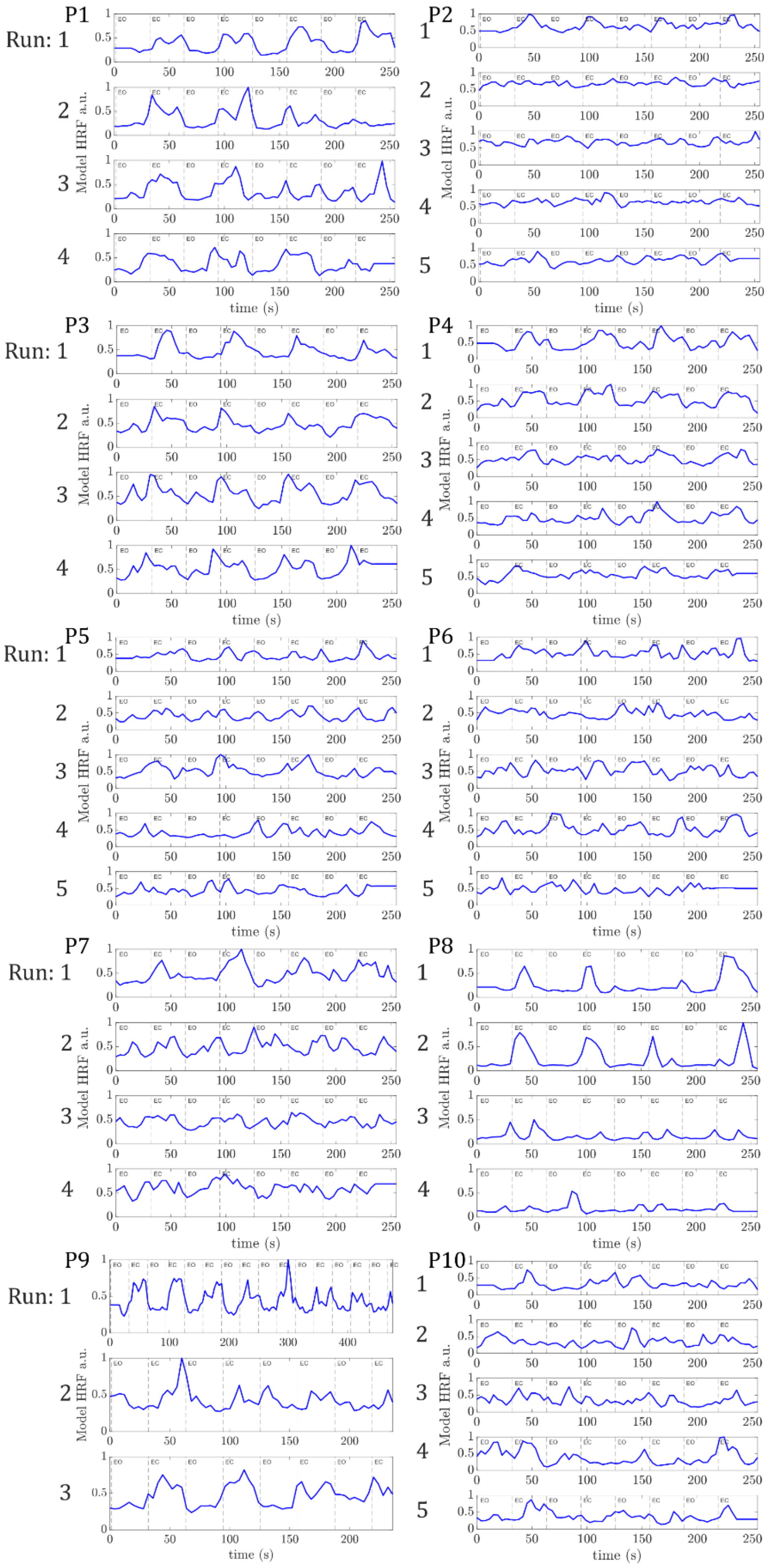


***Figure S1:*** *Regressors input to the GLM for each fMRI run shown for each participant. EEG alpha power VE time courses were convolved with the double-gamma HRF to create these regressors. Dashed lines denote periods of eyes open and eyes closed.*

### S2. Layer and column definition

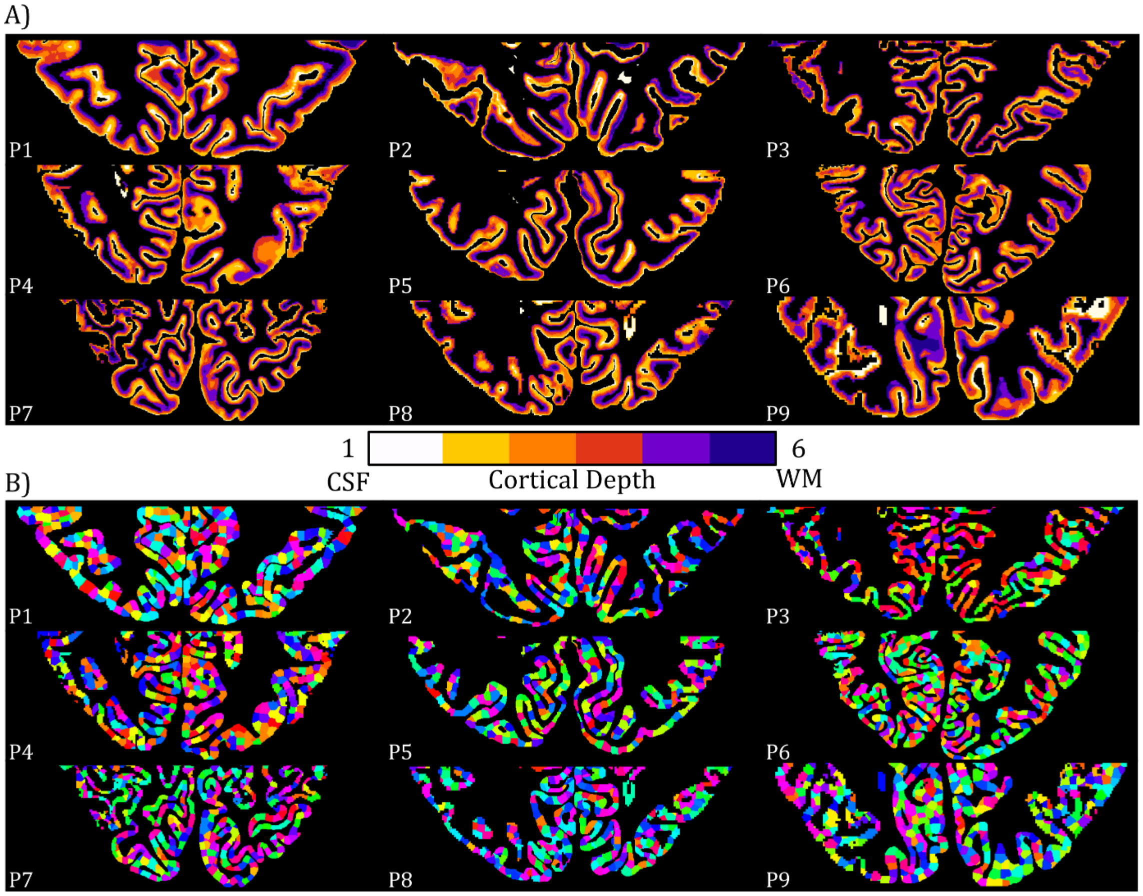


***Figure S2:*** *(A) The six equivolume cortical depths and (B) corresponding 4000 columns produced from LAYNII, representative slice shown for each participant. Colours in panel A represent the 6 cortical depths defined. Colours in panel B are random to show the parcellation of the cortex into columns.*

#### S2.1 Column Size

In the process of performing the analysis in Section 2.6.2 it became clear that not all of the small columns (when ROI was parcellated into 20000 columns) contain voxels in all layers. This was due to the geometry of the brain meaning that there may be no voxels defined in layer 1 or 6 over the very small area (i.e. 0.8-3.1 $mm^2$) of a column. The consequence of having no measured signal in a layer upon deveining was established. To mitigate this issue the cross-sectional area of the columns could be increased by reducing the number of columns. The effect of having 20000, 10000 and 4000 columns (corresponding to approximate column diameters of 1, 1.3 and 2 mm respectively) on the layer profile was assessed.

Figure S3 shows that the layer profile for the 'Uncorrected' data is a consistent shape for each column size. The 'Deveined' layer profiles also have a consistent shape, which only marginally changed with different numbers of columns. This data suggests that the profiles are not greatly affected by the size of the columns and therefore using the larger column size (smaller total number of columns) is preferable to maximise the presence of voxels (and therefore β-weights) in all layers which is more physiologically plausible. Therefore, remaining analysis was carried out using 4000 columns.

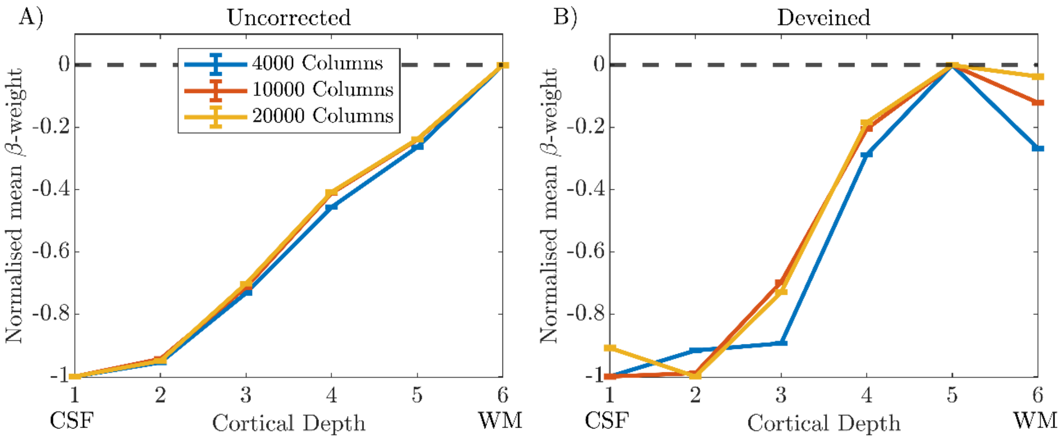


***Figure S3:*** *Layer profiles from V1 from Participant 1 shown for (A) 'Uncorrected' and (B) 'Deveined' data. Data is shown for 4,000 (blue), 10,000 (orange) and 20,000 (yellow) columns. Error bars show the standard error on the mean (SEM) for the mean normalised β-weight at each cortical depth.*

### S3. Choice of noise threshold for β-weights

To assess the effect of noise thresholds, three β-weight thresholds were assessed: i) none (all β-weights included), and excluding the ii) bottom 5% (i.e. keeping 95% of all voxels for deveining) and iii) bottom 10% (i.e. keeping 90% of all voxels for deveining) of absolute β-weights across all included columns. β-weights with an absolute magnitude below the threshold were set to "Not a Number" (NaN) and did not contribute to the deveining process or cortical depth profiles analysis.

Figure S4 shows the percentage of voxels within each 'included' column which contained positive and negative β-weights at different noise thresholds, with the visual cortex divided into 20000 columns. If no noise threshold was applied, there were a greater number of columns containing a high percentage of negative than positive β-weights, but still many columns with 10 – 50% positive β-weights (Fig S4B.i). Given the alpha-BOLD correlation is expected to result in negative β-weights in V1-V3, it was hypothesized that voxels with low amplitude, positive β-weights are dominated by noise.

The high percentage of positive β-weights within the unthresholded data reflect noise (β-weights close to zero) as is evidenced by the thresholding process. Applying the 5% threshold did not reduce the number of columns with a high percentage (>50%) of negative β-weights too greatly (Figure S4A.ii c.f. S4A.i); whilst the number of columns containing 10 – 50% positive β-weights reduced by a factor of 2.7 compared with no threshold for Participant 1 (Figure S54.ii c.f. S54.i). This was representative of the reduction factor over all participants 2.8±0.9 (mean±std), suggesting these β-weights below the 5% threshold were dominated by noise. Further increasing the threshold to 10%, resulted in nearly all data being within the lowest bin of the histogram for the positive β-weights (Figure S4B.iii), but this also led to an unacceptable reduction in the number of columns with a high percentage of negative β-weights (Figure S4A.iii), a pattern seen across all nine participants (data not shown). This suggests a 10% threshold was too stringent and so a 5% threshold was used.

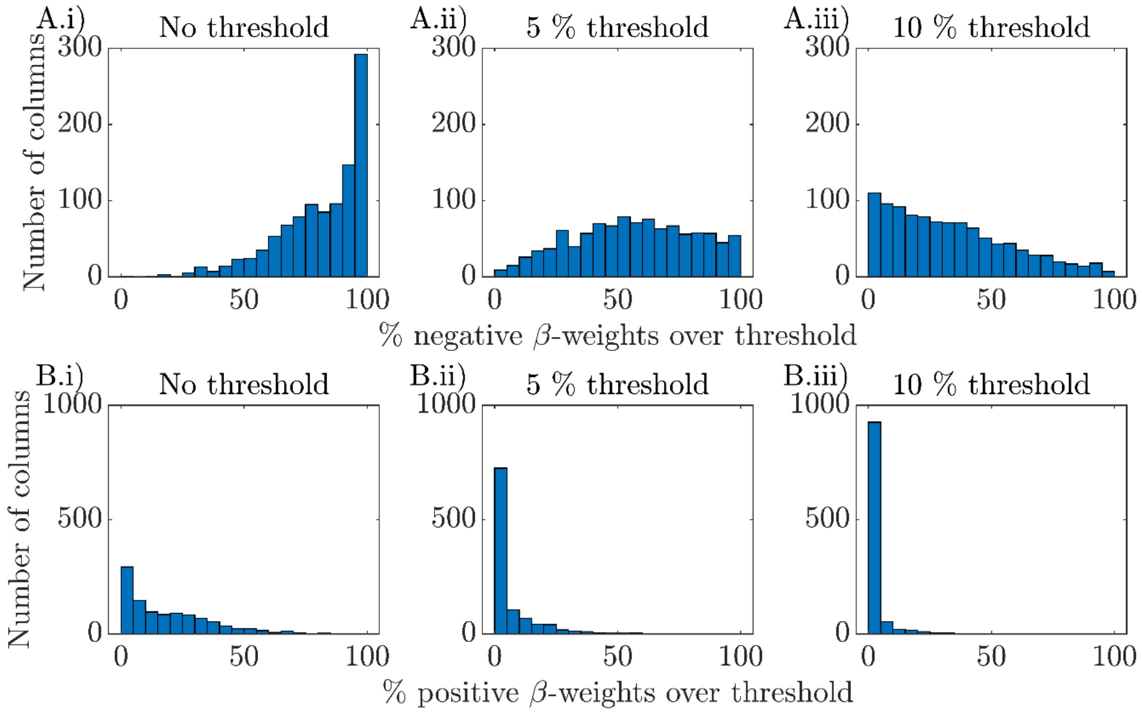


***Figure S4:*** *Histograms of the percentage of (A) negative β-weight voxels, and (B) positive β-weight voxels within columns across the three thresholding levels (i) no threshold (ii) 5% exclusion threshold and (iii) 10% exclusion threshold. Data shown for a representative participant (Participant 1).*

### S4. Vein Mask

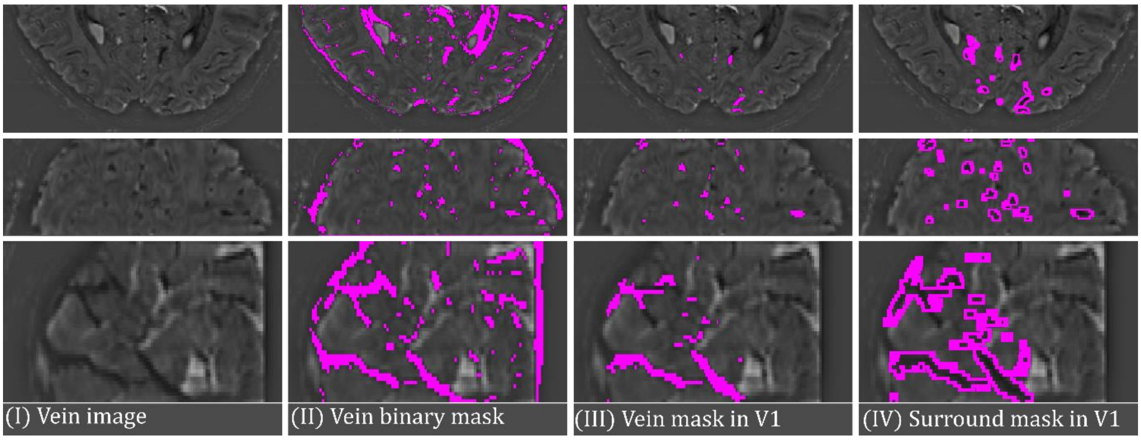


***Figure S5:*** *Generation of the vein mask in V1, with images shown in the axial (top), coronal (middle) and sagittal (bottom) view for a single participant. (I) Smoothed mean 3D-EPI image subtracted from unsmoothed 3D-EPI data (left column) to generate a 'Vein image', (II) 'Vein image' threshold in FSLeyes (middle column) to generate a binary mask which was (III) manually corrected to generate a vein mask in V1 (right column) (IV) the vein mask in V1 was dilated four-fold to generate the vein surround mask.*

### S5. Participant 10 GLM results

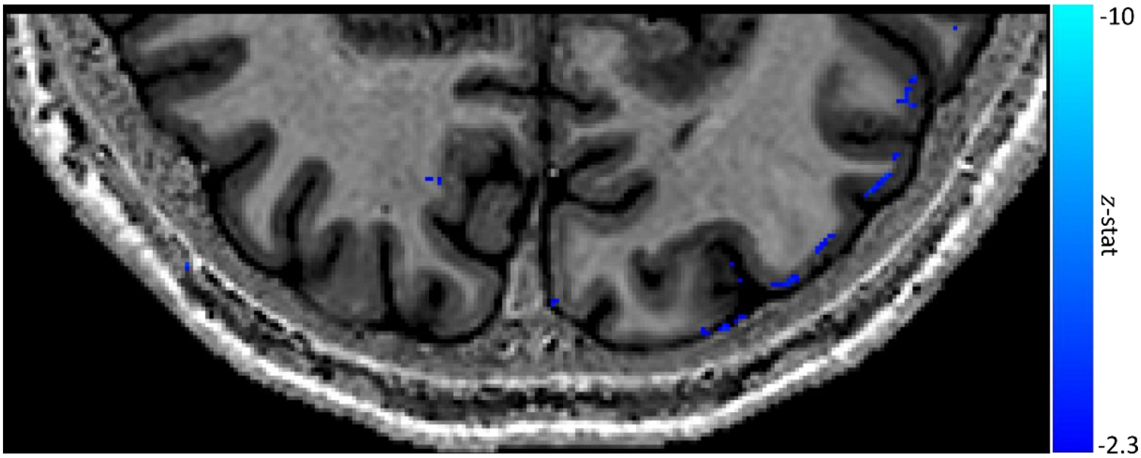


***Figure S6:*** *Negative contrast z-stat map (threshold at $z < -2.3$ and cluster corrected $p < 0.05$) from the fixed effects GLM using the EEG alpha power time course convolved with a double gamma HRF as a regressor overlaid on the PSIR for participant 10.*

### S6. Impact of regressors used in GLM

Figure S7 shows a comparison of the significant activation observed for each of the 3 GLMs performed: 1) GLM with regressor of only alpha power (plus nuisance regressors) 2) GLM with regressor of only boxcar denoting task (plus nuisance regressors) and 3) GLM with regressor of alpha power orthogonalised to boxcar (plus nuisance regressors). Areas of activation are generally the same when comparing between GLMs for each participant. As expected, the activation is weakest for the alpha power orthogonalized with the boxcar (GLM3) but the activation that is present matches the activation areas of the other two GLMs.

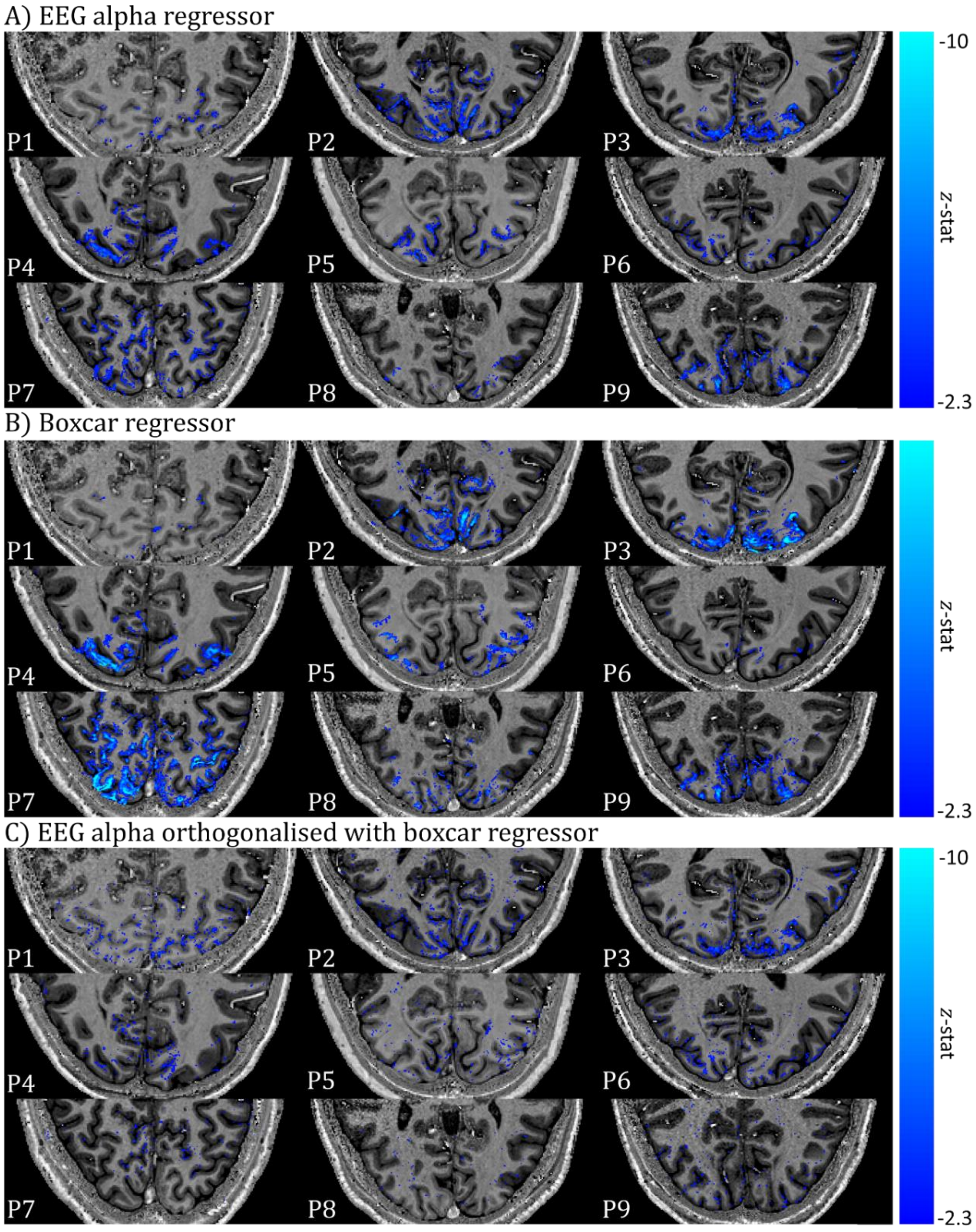


***Figure S7:*** *Comparison of negative contrast z-stat map showing regions of significant correlation for three different GLMs: A) EEG alpha power timecourse (as shown in Figure 6); B) a boxcar of eyes-open, eyes-closed periods; and C) EEG alpha power timecourse orthogonalized with the boxcar (which is also contained in the GLM). Fixed effects threshold at $z < -2.3$ and cluster corrected $p < 0.05$ are shown overlaid on each participant's PSIR. Note slices shown here are chosen to best compare across GLMs and are different to those shown in Figure 6.*

### S7. Effect of proximity to veins on layer profile

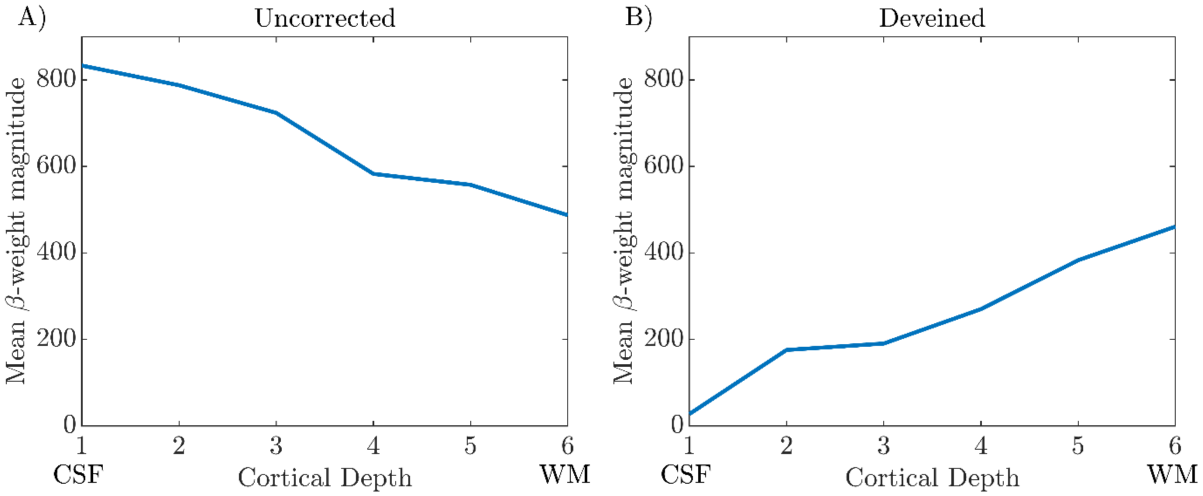


***Figure S8***: *Example from a single column from Participant 1 where the sign of the gradient of the cortical depth profile changes from negative to positive between the 'Uncorrected' (A) and 'Deveined' (B) data.*

### S8. Supplementary Tables

| Participant | EO | EC |
|---|---|---|
| P1 | 554 | 1121 |
| P2 | 274 | 295 |
| P3 | 314 | 517 |
| P4 | 258 | 424 |
| P5 | 478 | 631 |
| P6 | 525 | 538 |
| P7 | 309 | 397 |
| P8 | 343 | 705 |
| P9 | 271 | 601 |
| P10 | 344 | 441 |
| **mean±stdev** | **367±110** | **567±229** |

***Table S1***: *Mean alpha power from VE location for eyes open and eyes closed periods averaged over all runs for each participant.*

| Participant | No# z-stats $\times 10^5$ | V1 overlap % | V2 overlap % | V3 overlap % | GM overlap % | CSF dilated GM overlap % | WM dilated GM overlap % |
|---|---|---|---|---|---|---|---|
| P1 | 6.6 | 14.0 | 3.7 | 5.3 | 73.0 | 84.4 | 76.6 |
| P2 | 16.7 | 20.4 | 5.6 | 2.4 | 50.9 | 66.3 | 57.7 |
| P3 | 10.3 | 21.5 | 5.2 | 1.9 | 44.2 | 59.0 | 49.3 |
| P4 | 9.9 | 10.3 | 5.9 | 9.3 | 50.7 | 58.2 | 57.2 |
| P5 | 8.0 | 12.8 | 3.4 | 3.9 | 55.6 | 67.7 | 63.3 |
| P6 | 6.0 | 5.9 | 5.5 | 7.9 | 55.2 | 65.8 | 60.9 |
| P7 | 13.2 | 11.4 | 7.6 | 6.9 | 52.6 | 64.6 | 58.8 |
| P8 | 2.6 | 6.4 | 6.4 | 0.3 | 46.3 | 54.2 | 54.8 |
| P9 | 8.1 | 14.7 | 2.9 | 1.3 | 51.2 | 65.0 | 59.1 |
| **mean±stdev** | **9.1±4.1** | **13.0±5.4** | **5.1±1.5** | **4.4±3.2** | **53.3±8.2** | **65.0±8.6** | **59.7±7.4** |

***Table S2***: *Percentage overlap of the significant (z<-2.3) z-stat voxels with the GM ribbon and each of the visual ROIs (V1, V2 and V3), the entire GM ribbon and the GM ribbon dilated towards the CSF and WM separately.*
